# Resource: Scalable whole genome sequencing of 40,000 single cells identifies stochastic aneuploidies, genome replication states and clonal repertoires

**DOI:** 10.1101/411058

**Authors:** Emma Laks, Hans Zahn, Daniel Lai, Andrew McPherson, Adi Steif, Jazmine Brimhall, Justina Biele, Beixi Wang, Tehmina Masud, Diljot Grewal, Cydney Nielsen, Samantha Leung, Viktoria Bojilova, Maia Smith, Oleg Golovko, Steven Poon, Peter Eirew, Farhia Kabeer, Teresa Ruiz de Algara, So Ra Lee, M. Jafar Taghiyar, Curtis Huebner, Jessica Ngo, Tim Chan, Spencer Vatrt-Watts, Pascale Walters, Nafis Abrar, Sophia Chan, Matt Wiens, Lauren Martin, R. Wilder Scott, Michael T. Underhill, Elizabeth Chavez, Christian Steidl, Daniel Da Costa, Yusanne Ma, Robin J. N. Coope, Richard Corbett, Stephen Pleasance, Richard Moore, Andy J. Mungall, IMAXT Consortium, Marco A. Marra, Carl Hansen, Sohrab P. Shah, Samuel Aparicio

**Author notes:** equal contribution (Emma Laks, Hans Zahn, Daniel Lai, Andrew McPherson, Adi Steif).

## Abstract

Essential features of cancer tissue cellular heterogeneity such as negatively selected genome topologies, sub-clonal mutation patterns and genome replication states can only effectively be studied by sequencing single-cell genomes at scale and high fidelity. Using an amplification-free single-cell genome sequencing approach implemented on commodity hardware (DLP+) coupled with a cloud-based computational platform, we define a resource of 40,000 single-cell genomes characterized by their genome states, across a wide range of tissue types and conditions. We show that shallow sequencing across thousands of genomes permits reconstruction of clonal genomes to single nucleotide resolution through aggregation analysis of cells sharing higher order genome structure. From large-scale population analysis over thousands of cells, we identify rare cells exhibiting mitotic mis-segregation of whole chromosomes. We observe that tissue derived scWGS libraries exhibit lower rates of whole chromosome anueploidy than cell lines, and loss of p53 results in a shift in event type, but not overall prevalence in breast epithelium. Finally, we demonstrate that the replication states of genomes can be identified, allowing the number and proportion of replicating cells, as well as the chromosomal pattern of replication to be unambiguously identified in single-cell genome sequencing experiments. The combined annotated resource and approach provide a re-implementable large scale platform for studying lineages and tissue heterogeneity.

## Introduction

Resolving the molecular basis for dynamics of cell populations to single-cell resolution is a current frontier in tumour biology. In related fields this challenge has stimulated the development of many methods for single-cell transcriptional analysis (Macosko et al., 2015; Ziegenhain et al., 2017), however large scale analysis of single-cell genomes has lagged, in part because of the physical challenges of capturing DNA efficiently with even coverage across the genome (Gawad et al., 2016). Resolving structures of single cell genomes in tissues and cell populations is required for studying negative selection and rare cell populations in cancer and normal tissues. Resolution of genome replication states and sub-clonal mutational patterns also requires single genome sequencing analysis as bulk sequencing obscures signals from small subsets of cells. Several amplification based methods have been described (Navin et al., 2011; Zong et al., 2012; Hou et al., 2012; Ni et al., 2013; Gawad et al., 2014; Lohr et al., 2014; Wang et al., 2014; Baslan et al., 2012) however amplification introduces coverage bias and polymerase bias into sequences (Gawad et al., 2016) which can be difficult to account for and moreover, technical duplicate sequences cannot be easily identified.

Previously we introduced DLP (direct library preparation), a direct DNA transposition single-cell library preparation method in nanolitre volumes that requires no pre-amplification, reducing the biases of the method compared to pre-amplification based approaches (Zahn et al., 2017). Nanolitre microfluidic reaction chambers not only made this method more economical compared to microlitre approaches, it allowed adaptation of DNA transposase chemistry to a single-cell template directly without the need for whole genome amplification (WGA). Various other WGA based methods of single-cell whole genome sequencing (WGS), (Navin et al., 2011; Zong et al., 2012; Hou et al., 2012; Ni et al., 2013; Gawad et al., 2014; Lohr et al., 2014; Wang et al., 2014; Baslan et al., 2012) including degenerate oligonucleotide-primed PCR (DOP-PCR), multiple displacement amplification (MDA) and multiple annealing- and looping-based amplification cycles (MALBAC), are affected by increased bias or polymerase errors. DLP was previously shown to outperform DOP-PCR for coverage uniformity. The recently introduced single-cell combinatorial indexed sequencing (SCI-seq) aims to increase the throughput of single-cell sequencing but is limited by its high duplication rate and relatively low coverage breadth (Vitak et al., 2017). Despite the performance of microfluidic-based DLP analysis, the use of custom microfluidic devices presents an obstacle to adoption in many labs and also places limits on scalability due to fabrication constraints. They also can impose constraints on cell size, with large cells clogging channels, and very small cells passing through traps, unless devices are customized for the cell type. Similar limitations on cell size apply to some droplet based methods.

To address these limitations, we have developed and optimized an alternative and much higher-throughput implementation of nanolitre volume DLP, referred to here as DLP+, that is based on high-density nanowell arrays and picolitre volume piezo dispensing technology available “off the shelf” (Figure 1). To achieve comparable performance to our microfluidic approach, we rigorously optimized key reaction parameters, thus providing insight into determinant conditions for high fidelity sequencing of single genomes across a wide range of tissues and cells. We show that optimized DLP+ enables robust and scaled analysis of tens of thousands of cells per experiment across various tissue types. We couple the experimental work flow with a required cloud-enabled computational platform which performs scalable and automated quality control, copy-number segmentation of thousands of low-coverage genomes and an interactive data visualization interface for real time browsing and filtering of data. Using this resource, we show how clonal merging of complex genomes can be used to recover features to single nucleotide resolution and illustrate how rare mutational processes and genome states can be studied in tissues using these approaches.

**Figure 1.**
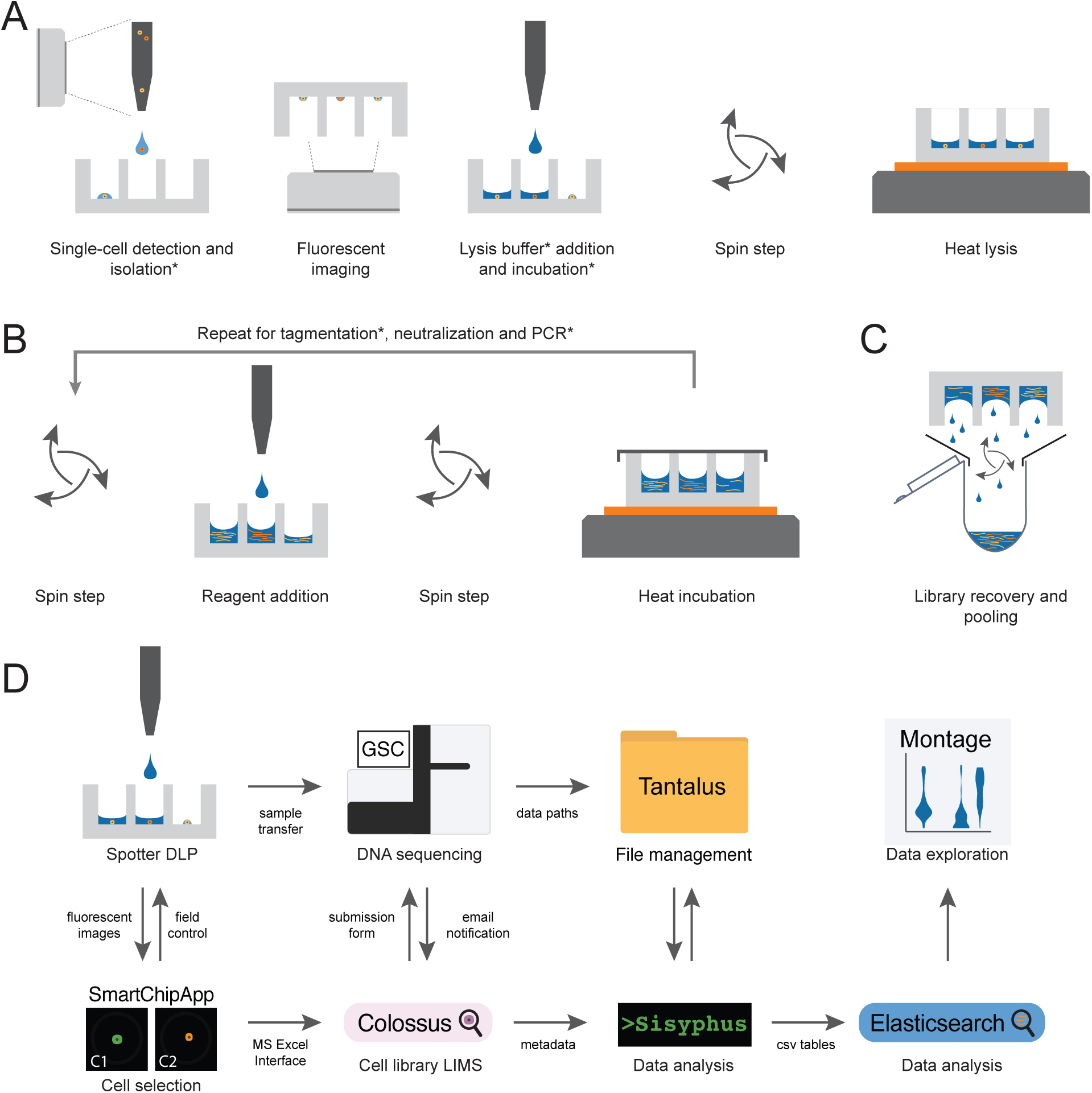
Concept schematic of the experimental and computational processes for DLP+. **a** Cell isolation and lysis. A cell suspension is loaded into a dispensing nozzle; single cells are identified using real-time imaging during dispensing and selected cells are isolated in individual nanowells preprinted with unique duel index sequencing primer pairs; occupancy and cell state are confirmed by fluorescent imaging and recorded; wells can be selectively addressed to add cell lysis solution; spin and seal steps are used to collect reagents in the bottom of the well and prevent evaporation during heat lysis. b Open-array library construction. DLP+ libraries from unamplified single cells are built by carrying the chip through a series of reagent addition, spin, seal, and heat incubation steps. c Pooled recovery. Indexed single-cell libraries are pooled for multiplexed sequencing. d Pipeline workflow for single-cell genome data management, alignment and post processing. Sample data is entered into a custom single-cell library LIMS, Colossus. After scanning and processing the fluorescent images of isolated single cells in chip using the custom SmartChipApp, details on the single cells including cell state are uploaded to Colossus for single-cell library tracking as well as being used for spotting robot control to individually address and lyse selected cells. After library construction Colossus automatically generates a submission form and the pooled single-cell library is passed to the Genome Science Centre (GSC) for sequencing. The GSC sends an email notification when sequencing is completed and the data is transferred by Tantalus, a file management system, and analyzed by Sisyphus, a workflow automation pipeline. CSV tables output by Sisyphus are loaded into Elasticsearch and visualized with a custom interactive data visualization tool, Montage. *These steps were optimized to ensure DLP performance in nanowells.

## Results

### A resource of diverse, annotated, single cell genomes generated with scalable open array transposon mediated DNA sequencing

#### Scalable single-cell library preparation in open nanolitre wells

To scale amplification free transposon based single-cell genome sequencing to thousands of cells per library we implemented DLP+ using commodity off the shelf core components, specifically a contactless piezoelectric dispenser (sciFLEXARRAYER S3, Scienion) and open nanowell arrays (SmartChip, TakaraBio; Figure 1, Figure S1a). In the approach described here each chip contains 5184 wells (72×72 array) with a volume of 110nL per well and several chips can be processed in a day with one instrument. Key to implementation of the molecular steps for genome sequencing library construction, the spotting robot can address and fill each well in increments of 300-550 pL (Figure S1b) thus providing approximately 200-360 freely programmable reagent addition steps. A further important feature is the use of real-time object recognition (Figure 1a, Figure S1c-e) during piezo droplet dispensing through an on board camera (cellenONE, Cellenion, further details in methods). This allows for control of the cell spotting pattern on open arrays and also permits selection of cells from debris and other objects, bypassing the limitations of Poisson dilution cell loading (Figure S1f-g). The details of the platform are described in Figure S1.

For cell indexing, dual sequencing index primers are pre-spotted (Figure S2a) to label each chamber for pooled recovery. This provides a more streamlined library preparation protocol, since the same PCR reagent mix can be delivered to all wells in parallel. Since imaging occurs before the library preparation reagents are spotted, doublets, empty wells, or cells with contamination can be excluded from library preparation (Figure 1a). The final libraries are pooled during recovery (Figure 1c) and sequenced at the desired coverage depth on HiSeq instruments (Illumina). After the libraries are de-multiplexed, aligned, and analyzed for copy number alterations (CNA), single nucleotide variants (SNVs) and other features as described below, the processed single-cell data are loaded into a custom visualization platform. Below, we describe the key reaction attributes and working ranges for single-cell genome library construction using this scaling approach.

#### A scalable cloud computing pipeline for single-cell genome analytics

Analysis of DLP+ single-cell whole genome data requires alignment of sequence reads to the genome reference, and further processing to execute normalization, quality control, genome copy number analysis and data visualization. This is a computationally non-trivial task, considering that a typical single library of 1000-3000 single-cell genomes, requires computing over hundreds of thousands of files and processes.

We developed a scalable cloud-computing, open source software platform for automated analysis of DLP+ data (Figure S4a). The platform is comprised of three key components (i) Tantalus, a relational database implementation for tracking metadata and file information relating to each single cell, (ii) Sisyphus, a computational pipeline coupled to the database for triggering whole genome alignment, post alignment quality assessment and copy number inference including ploidy estimation, (iii) Montage, a web-based interactive data visualization platform for real-time browsing of the derived results (Figure 1d) on single cells and over groups of cells. Montage supports creation and sharing of collections of visualizations-a dashboard-and is equipped with features to facilitate data exploration and quality assessment: views are interactively linked such that a selection in one view is reflected in all others, and researchers can dynamically filter plotted data or alter the plot dimensions (Figure S5). Montage provides standard visualization types, such as a scatterplot (Figure S5f) over key pre-computed quality control attributes, but additionally includes custom views to enable the exploration of thousands of single-cell copy number profiles (Figure S5c).

The supplement contains a use case exemplifying Montage QC dashboard for inspecting libraries, supported by a read-only version of Montage for public browsing and downloading of all the post-alignment single-cell data, available on publication at https://www.cellmine.org. This version contains unfiltered data and preserves all the interactive view linking and dynamic filtering for data exploration.

#### Distinguishing library quality from biological states with a feature-based classifier

We observed that poor quality libraries generate sequence and alignment properties that are distinct from biological states (Figure 2a, b) and can be easily distinguished by inspection. However, manual inspection is neither scalable nor quantitatively defined for reproducibility, and thus we developed a machine learning classifier to automate quality control. We labeled ∼20,000 genomes manually through inspection as low/high quality assumed to represent failed and successful libraries. We computed 18 cell-specific features quantitatively capturing alignment and copy number metrics and trained a binary random forest classifier (Methods) using the labels as supervised classes. The trained model classifies new data, emitting the probability of a successful library given the 18 features. We refer to this probability as a quality score (QS), ranging from 0 to 1 (0.0 = failed, 1.0=perfect). The QS although continuous, tends to be bimodal (Figure 2c). For the present resource we established by calibration against previously published DLP (Zahn et al., 2017) that QS ≥ 0.75 efficiently separates poorly performing libraries from high quality libraries.

**Figure 2.**
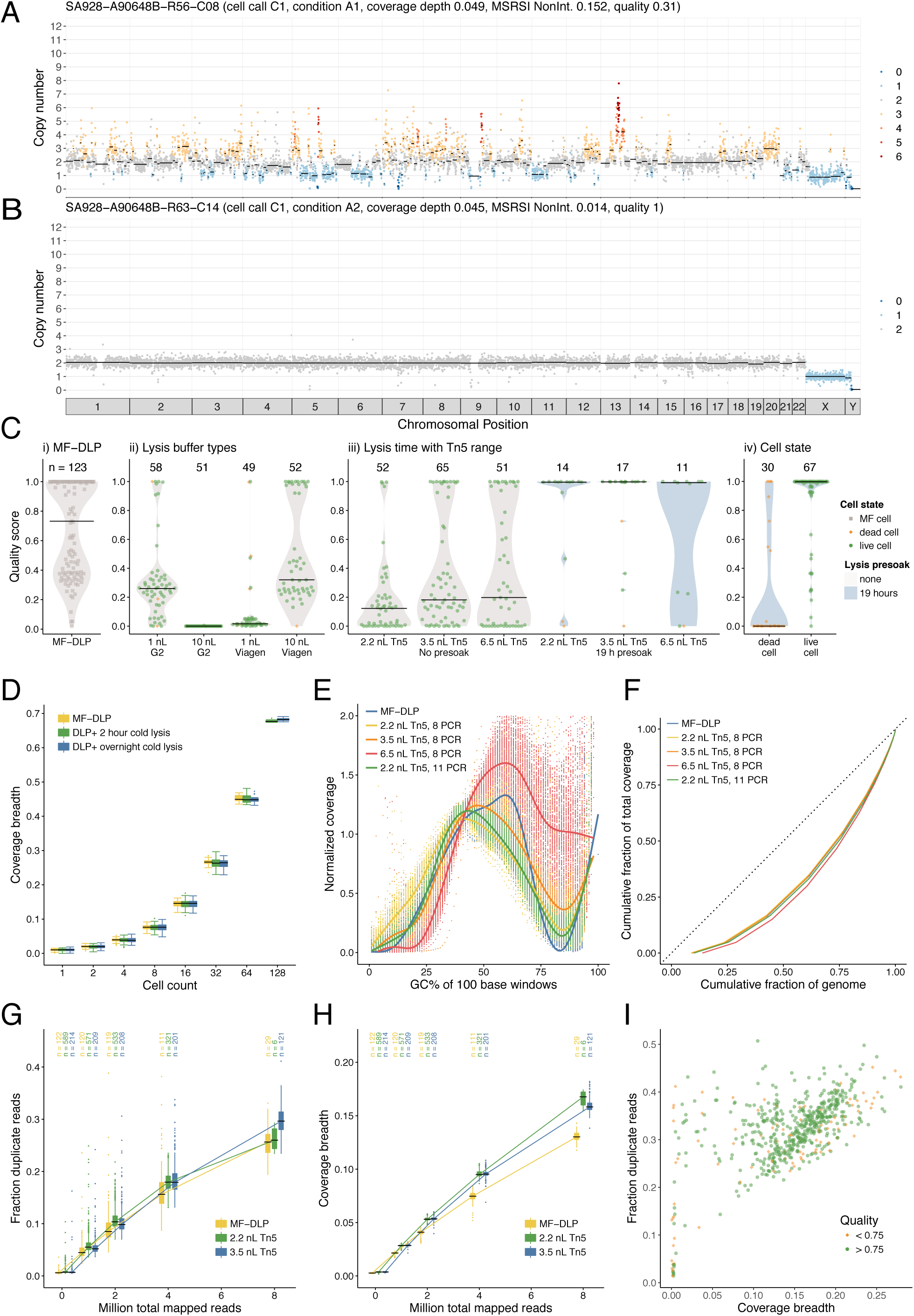
Optimization of DLP+ scWGS library construction for the open-array format. Examples of a high quality and b poor quality single-cell genome libraries from a diploid GM18507 lymphoblastoid (male) cell line. Colours correspond to integer HMM copy-number states (Ha et al., 2012); black lines indicate segment medians. c Quality score distribution over GM18507 cells of (i) the original MF-DLP data (Zahn et al., 2017)), (ii) lysis buffer types, (iii) Tn5 concentrations and increased lysis presoak times and (iv) cell state (live or dead). Numbers of cells are indicated above each violin plot, where black lines show medians and dots indicate individual cells (green circle = live, orange diamond = dead, grey square = no cell state data). Grey background indicates where cells underwent heat lysis immediately after lysis buffer addition, and blue background indicates cells kept in lysis buffer for 19 hours at 4 °C before heat lysis. d Distribution of coverage breadth of bootstrap sampling of GM18507 libraries using a 2 hours and overnight presoak lysis compared to a microfluidic device (MF-DLP (n = 122, (Zahn et al., 2017)), DLP+ 2 hours (n = 148), DLP+ overnight (n = 133). e GC bias of GM18507 libraries as a function of Tn5 concentrations and 8 or 11 PCR amplification cycles. f Lorenz curves showing genome-wide coverage uniformity of merged single-cell libraries over Tn5 concentrations and 8 or 11 PCR amplification cycles (downsampled to 64 cells per experimental condition). Dotted straight black line indicates perfectly uniform genome. g Distribution of fraction duplicate reads for GM18507 cells (2.2 nl Tn5, n = 587 (green); 3.5 nl Tn5, n = 571 (blue)) and on a microfluidic device (n = 141, (Zahn et al., 2017) (yellow)). The top column labels state the numbers of cells per condition. h Coverage breadth for the downsampling analysis of the same libraries depicted in panel g. i Fraction duplicate reads vs coverage breadth of deeply sequenced GM18507 libraries (3.5 nl Tn5, n = 571), 10 HiSeqX lanes) with low quality (<0.75) and high quality (>=0.75) indicated.

#### Biological and physical determinants of high quality DLP+ library construction

We initially applied the same reaction conditions from microfluidic DLP (Zahn et al., 2017) to establish amplification-free single cell WGS in open arrays (DLP+). However this resulted in many poor quality libraries measured by a high proportion of: i) alignments for which interpretable, integer state copy number profiles could not be inferred and ii) failed libraries where coverage was low or absent (Figure 2a, c ii: 1 nL G2 buffer). We therefore sought to quantitatively establish the physical reaction determinants of high quality libraries (e.g. Figure S4b), based on the computed quality score from the classifier. We systematically varied and evaluated several factors: cell lysis volume and buffer type; transposase (Tn5) concentration; post-indexing PCR cycles; cell lysis/DNA solubilisation time; and cell viability state.

We observed the following properties as determinants of high quality libraries. Cell lysis volume and buffer type exhibited a combinatorial effect, whereby increased volumes to avoid meniscus effects and evaporation required specific buffers that could be diluted in one pot reactions without impacting subsequent reactions (Figure 2c ii). As expected, increasing transposase (Tn5) concentrations increased library success (Figure 2c iii), but with a tradeoff of increasing bias in sequence GC representation (Figure 2e) and consequently genome coverage (Figure 2f). Similarly, we observed that an increase in post-indexing PCR cycles also tended to increase GC-bias (Figure 2e, f). The optimal value for most cell types was found to be approximately twice the Tn5 concentration of previously published microfluidic DLP (MF-DLP), with 8 post transposition cell indexing PCR cycles. This resulted in GC profiles and coverage very similar to MF-DLP and bulk libraries. Furthermore, cell lysis/DNA solubilisation time after cell isolation was a key determinant of high quality libraries. We established that incubating cells in lysis buffers for >2 hours was critical to robust solubilisation/transposase access of the genome (Figure 2c iii, d). Finally, the cell viability state (live or dead, determined by dye exclusion) impacted overall success as might be expected (Figure 2c iv), however some proportion of cells labelled as dead nevertheless produce high sequence quality libraries.

We next investigated how the increased lysis time affects the coverage breadth of merged single-cell libraries to produce high-quality bulk-equivalent genomes. We carried out bootstrap sampling and merging analysis of GM18507 single-cell libraries prepared using the 2 hours and overnight lysis protocols and compared the results with single-cell libraries prepared from the same cell line using our previously published MF-DLP protocol (Zahn et al., 2017) (Figure 2d, Figure S2b). To provide a meaningful comparison, we only used live cells, removed duplicates and uniformly downsampled each single-cell library to approximately 0.01X mean coverage depth (DLP+ 2 hours to 72.7%, DLP+ overnight to 88.9%, and MF-DLP (Zahn et al., 2017) to 8.2% of unique non-duplicate reads). Bootstrap sampling (1-8 cells n = 360 draws, 16 cells n = 240 draws, 32 cells n = 120 draws, 64 cells n = 60 draws, 128 cells n = 40 draws) and merging of 64 single-cell libraries resulted in a median coverage breadth of 43.7%, 44.4%, 43.3% in the MF-DLP, DLP+ 2 hours and DLP+ overnight libraries, respectively. Given that differences in coverage breadth between the lysis conditions were not significant (KW test: p = 0.6766), we determined that the longer lysis condition was preferable, since it permitted the protocol to be split between two reasonable-length work days.

We then evaluated optimal and non-optimal reaction parameters relative to the microfluidic GM18507 DLP dataset (Zahn et al., 2017), using bootstrapped subsampling of libraries to compare performance at similar read depths. It should be noted that the MF-DLP dataset was previously shown to produce equivalent breadth and uniformity to that of a bulk genome at the same coverage depth (Zahn et al., 2017), establishing a reasonable benchmark for comparison. Before bootstrap analysis, all single-cell libraries were downsampled to equivalent mean coverage depth of 0.05X (KW test, p= 0.16). Merging 64 downsampled genomes resulted in 3.16X median coverage depth and 90% median coverage breadth across all libraries (Figure 2f). However, the 6.5nL/8PCR condition and the 2.2 nL/11PCR condition had a significant lower genome coverage breadth (86.2% and 89.9%, respectively) compared with the MF-DLP dataset and both the 2.2nL/8PCR and 3.5nL/8PCR conditions (90.6%, 91.3%, 90.7%, respectively; KW test,p= 1.542e-13). In fact all contrasts between MF-DLP, 2.2nL/8PCR, nL/8PCR and the 6.5nL/8PCR, 2.2nL/11PCR condition were significant (post hoc Dunn’s test with Benjamini-Hochberg correction), but there was no significant difference between MF-DLP, 2.2nL/8PCR, 3.5nL/8PCR conditions; see Table S 3) at the same coverage depth (KW test, p = 0.6974), indicating this to be the optimal conditions/parameters. Pooling 128 cells at a mean coverage depth of 0.05X per cell resulted in 96.9% coverage breadth at an aggregated depth of 6.35X for the 2.2nL/8PCR protocol. In comparison, the 6.5nL/8PCR condition achieved significantly less genome coverage (93.2%) at the same depth (Table S 3, KW test p = 0.3302). This suggests that higher GC-bias associated with increased Tn5 concentration indeed reduced genomic breadth. In addition, through Lorenz curve analysis, we found that the 6.5 nL Tn5 condition was considerably biased, while there was no major difference between the MF-DLP dataset and the 2.2nL and 3.5 nL Tn5 conditions (Figure 2f, Figure S2c). For the 2.2nL and 3.5 nL Tn5 conditions, 64 merged single cells achieved a comparable coverage breadth and uniformity as 64 merged single-cell genomes from the microfluidic dataset (Figure 2f). We then compared genome coverage breadth, sequence reads and duplicate reads and established that under optimal conditions DLP+ performs very closely to MF-DLP (Figure 2 g, h, Figure S2g). Moreover the rate of duplicate accumulation with increasing depth indicates acceptable rates of between 20-40% duplicates even at ∼0.2 genome coverage (Figure 2i). We note that at ∼0.1 genome coverage ∼48-64 cells are sufficient to reconstruct a bulk equivalent genome.

Thus, we established an optimal parameter range for high quality, scalable single cell genome sequencing, without template amplification, on open arrays and with an off the shelf picolitre dispensing robot. These advances substantially increase throughput over the MF-DLP method from hundreds to thousands of cells, and improve overall quality of the libraries relative to the original method (Figure 2c i).

We next applied optimized DLP+ across a range of different tissue and cell types including cell lines, human breast cancer patient derived xenograft (PDX) samples, and tumour samples from mouse model of synovial sarcoma (SS) and human follicular lymphoma (FL), thereby generating a reference resource of high quality annotated single-cell genomes (Figure 3, Table S 2). Cells from these samples were chosen to cover a range of cell sizes from 5 microns to 80 microns and included cells fresh from culture (cell lines), cells isolated from cryopreserved tissues (breast PDX, FL), and cells from dissociated primary tumour material. We then applied the multi-parameter QS classifier to 41270 single cells and nuclei, establishing a probabilistic quality score (QS) on each cell. We generated libraries from 41270 single cells and using a QS cut-off value of 0.75 to filter out low coverage and failed cells, we observed that 70.0% of cells identified as live (n = 22815/32576) and 30.7% of cells labelled as dead (n = 2261/7259) produced high quality single cell genome sequences ((Figure 3a,b). Of tissue and cell line samples where live and dead cells were included (n = 30), 90% had a significant difference in quality between live and dead cell (linear regression, p > 0.05, Table S 3). The DLP+ process allows for dead cells to be selectively excluded from library construction based on their fluorescent staining, however we chose to include them, in order to evaluate the effect of cell viability on successful library construction in different tissue types and provide a full reference set of genomes in differing biological states. Dead cells with QS ≤ 0.75 had a very low median per cell read count of 19740, rather than resulting in poor quality libraries with representation bias and non-integer copy states. We therefore included cells labelled as dead or dying by vital dye staining, but which nevertheless produce high QS libraries.

**Figure 3.**
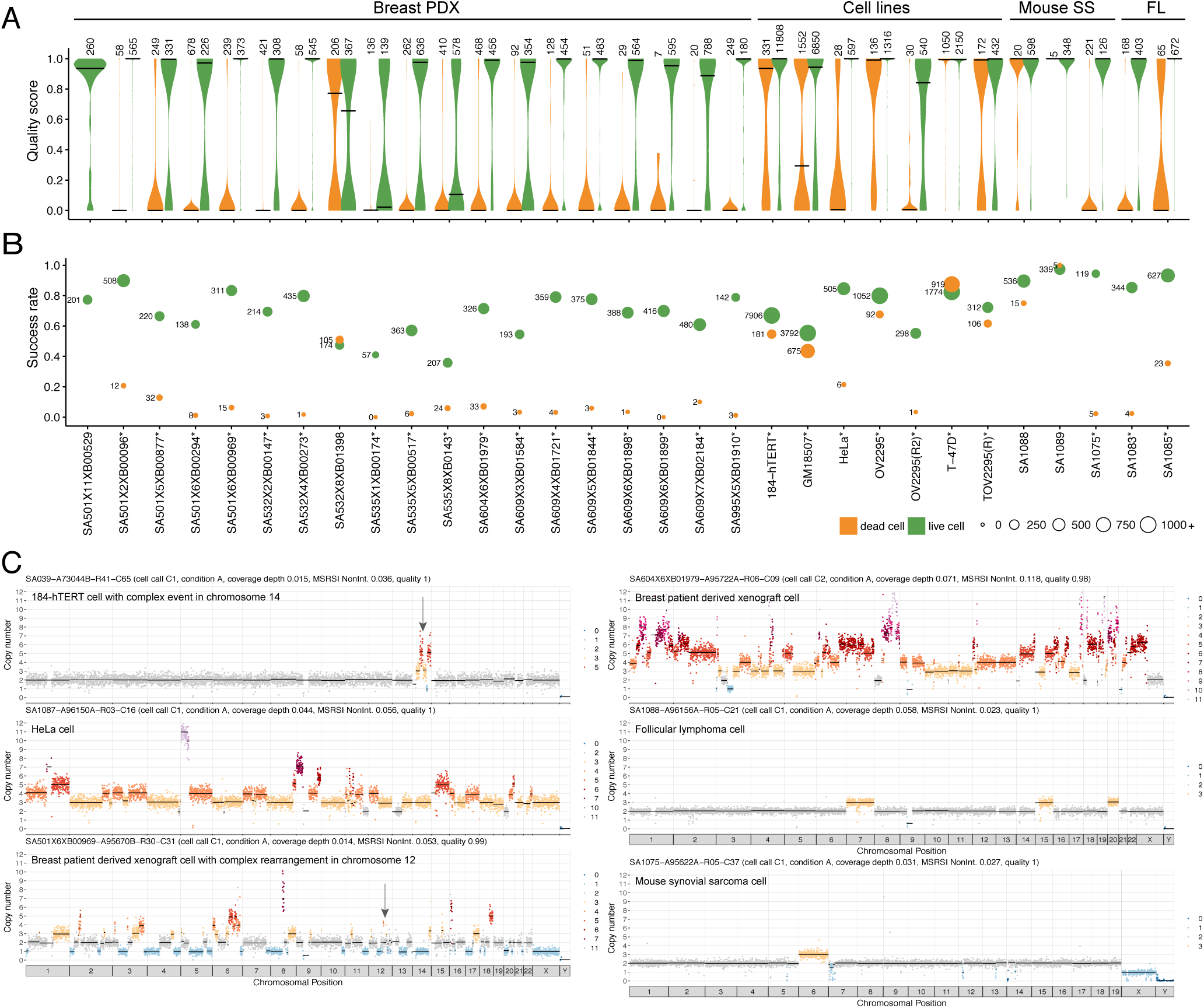
caption below Figure 3 DLP across different tissue types split by viability: live cells (n = 32576, green) dead cells (n = 7259, orange). **a** Violin plots showing the quality score of single-cell libraries across various tissue types, split by cell viability status (live or dead), with number of cells shown above the violin. Black lines show median. b Fraction of successful cells in a sample (quality >0.75), split by cell viability. The size of the bubble represents the total number of successful cells. Violin and bubble colours indicate cell viability. Samples where the difference in quality of all of the live and dead cells had a p value of >0.05 (linear regression) are indicated by an asterisk. c Example single-cell copy number profiles from cell lines, breast PDX, follicular lymphoma, and mouse synovial sarcoma. Colours correspond to integer HMM copy-number states; black lines indicate segment medians.

Key to flexibility of DLP+ for tissue is the ability to handle nuclei as well as cells, we observed that 69.9% of nuclei had QS above 0.75 (n = 933/1335), showing that DLP+ is equally successful producing high quality libraries from live cells or nuclei (Figure S3). Nuclei and cell CN profiles from the same sample cluster together rather than forming two clusters (Figure S3b) and had similar QS, total mapped reads, duplicate reads, and integerness within libraries where cells and nuclei were prepared in parallel (Figure S3c), producing copy number profiles of equal quality (Figure S3d).

Across different tissue/cell types, DLP+ produced equal QS distributions on live cells (KW test, p = 5.91E-238, Table S 3), demonstrating the flexibility of the method on both tissue types, and on samples that have simple diploid (SS, FL) or complex and sometimes polyploid genomes (breast PDX) (Figure 3a, c). The rate of library construction success (QS > 0.75) for live cells ranged from 35.4% to 97.4% depending on the sample (Figure 3b), with 28/31 libraries having a live cell success rate over 50%, median 72% success rate over all libraries. Notably, QS>0.75 DLP+ libraries can identify highly aneuploid genome states, including complex rearrangements (Figure 3c in single cells, in a similar manner to DLP. As noted below, merging of cells into clones permits resolution of these features to single nucleotide resolution without prohibitively sequencing to depth every cell. Taken together, the data illustrate the scalability and broad applicability of DLP+ to single cell genome sequencing.

### Merging cell sub-sets with shared copy number enables inference of clone-specific single nucleotide resolution events and clonal phylogenies

Single-cell sequencing techniques promise to provide more accurate measurement of clonal genotypes and clone proportions in cancer samples owing to direct demultiplexing from DNA barcodes, and thereby obviating cumbersome and error-prone bulk tissue computational deconvolution methods. This in turn facilitates accurate phylogenetic reconstruction of major clones in a cancer. An important practical tradeoff is the coverage per cell of genome sequenced against the number of cells from a population. We hypothesized that large numbers of DLP+ single cell genomes sequenced at low coverage, could be leveraged to determine clonal populations and subsequently infer clone-specific nucleotide resolution somatic events including SNVs, allele-specific copy number and rearrangement breakpoints plus phylogenetic trees computed over these events. To address this, we developed an analytic workflow (diagrammed in Figure S11) which first predicted germline and somatic SNVs, and breakpoints on a merged dataset where all cells were collapsed into a ‘pseudobulk’ genome. We then clustered single cells into cell subsets by their copy number profiles and measured the presence/absence of somatic and germline SNVs, and rearrangement breakpoints in each clone. Given measurements of variants per clone, we then calculated allele specific copy number per clone and inferred phylogenetic evolutionary histories given SNVs, breakpoints and copy number profiles.

To exemplify this approach, we generated 1966 DLP+ libraries from 3 clonally related high grade serous (HGS) ovarian cancer cell lines derived from the same patient, sourced from one primary tumour and two relapse specimens. On cells with > 500,000 reads (n = 1542 cells retained) and quality score > 0.5 (n = 1345 cells retained) we used dimensionality reduction and clustering (Supplemental methods) to identify 9 cell sub-sets with shared copy number profiles as a first approximation to clones (subsets with >=50 cells, 891 cells retained). The 9 clones ranged in size from 62 to 145 cells with an average coverage depth of 5X Figure S6.

For each clone, we computed clone-specific features including total copy number ((Figure 4a), allele-specific copy number, SNVs and breakpoints. For allele specific copy number, we inferred haplotype blocks from germline polymorphisms using Shape-IT (Delaneau et al., 2011) and the 1000 Genomes phase 2 reference panel. Across the 9 clones, we obtained an average of 91 to 222 reads per haplotype block, and an average of 21,219 to 30,941 haplotype blocks with 100 or more reads. Clone-specific haplotype block allele ratios coincided with fractional values that could easily be matched to genotypes consistent with clone specific copy number calls. For each clone, we thus fit a straightforward HMM to infer minor copy number based on haplotype block read counts and total copy number (Figure 4b). By way of example, total copy number and minor allele fraction for clone 0 (Figure 4a,b, 145 cells) is consistent with a whole genome duplication (WGD) event. Chromosomes 1,7,10,11 all harbour 4 copies and a minor allele fraction near 0.5. Furthermore chromosomes 2, 5, and 9 all contain segments with 3 copies and minor allele fraction of 0.33. These events are consistent with single copy loss from a WGD event. By contrast, chromosomes 3,4,6 and 12 all harbour segments with 5 copies and minor allele fraction of 0.4 consistent with a single copy gain (e.g. AAABB) after WGD. Additionally, DLP+ allows for the resolution of clone specific focal amplifications such as the 4 copy segment of chromosome 13 specific to the clone 1, 8 branch of the phylogeny (Figure S9, an event that would be difficult to characterize from merged data of OV2295 (Figure S7). Finally, we interpreted segments with minor allele fraction of 0 as loss of heterozygosity events. These are evident directly from the data, for example chromosome 17, known to be homozygous in nearly 100% of HGS ovarian cancers, is unambiguously centered at 0 minor allele fraction.

**Figure 4.**
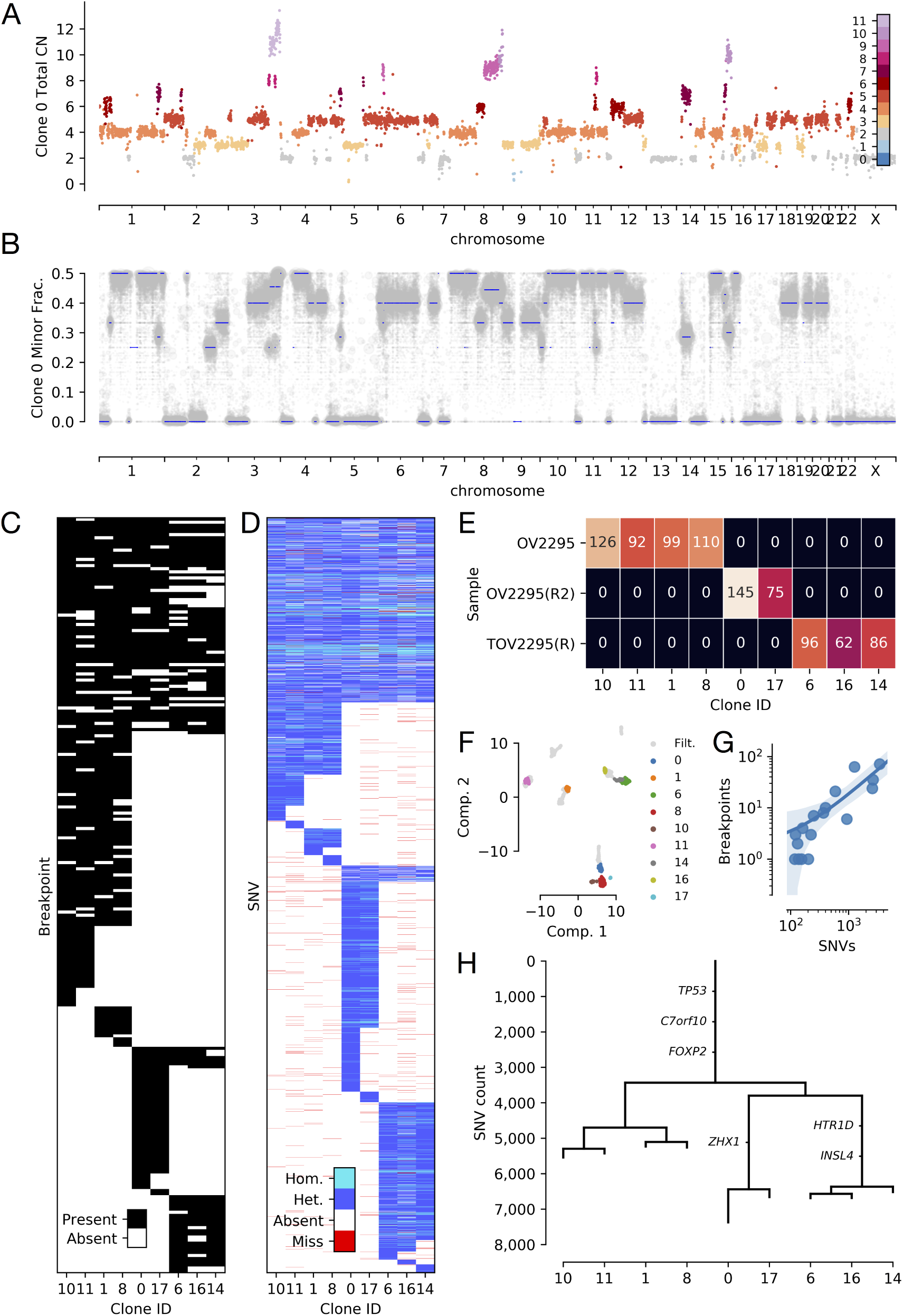
Features from merging of clones of OV2295, OV2295(R2), TOV2295(R) cell lines based on single-cell CNV (n = 891). **a** Raw total copy number for clone 0 (y-axis) across the genome (x-axis) colored by inferred total copy number. b Minor allele frequency of clone 0 (y-axis) across the genome (x-axis) with inferred minor copy number ratio (minor copy number / total copy number) shown as blue lines. c Presence of breakpoints (y-axis) in each clone (x-axis). d Presence and state of SNVs (y-axis) in each clone (x-axis) with SNVs with no coverage in a clone shown in red, heterozygous and homozygous SNVs as determined by reference and alternate allele counts shown in dark and light blue respectively. e Cell counts per clone per sample. f Reduced dimensionality representation of n=1345 cells passing preliminary filtering, with cells excluded by additional filtering in grey, as calculated using UMAP. g Correlation between counts of breakpoints and SNVs on the branches of the identically structured phylogeny inferred for both variant types. h Phylogenetic tree with branch lengths calculated as counts of SNVs originating on each branch.

We further explored the inferred clusters using both SNVs and breakpoints. We used mutationSeq (Ding et al., 2012) and Strelka (Saunders et al., 2012) to identify 13697 SNVs across the 9 clones, and maximum likelihood to infer a phylogenetic tree relating the inferred clones (Figure 4d, h). As expected, each of the 3 samples formed a distinct clade in the phylogeny. A total of 11078 SNVs fit perfectly with the inferred phylogenetic tree with 3352 predicted as ancestral, 2469 clone specific and the remaining 7876 clade specific. Ancestral mutations with significant impact include TP53 (584T>C), FOXP2, and C7orf10. Clade specific mutations include ZHX1, HTRD1 and INSL4. We then used an orthogonal method (hierarchical clustering) to infer a phylogeny from breakpoints inferred using deStruct (McPherson et al., 2017a) (Figure 4c). A total of 169 of the 244 total breakpoints fit perfectly with the inferred phylogeny. By maximum parsimony, 70 breakpoints were predicted ancestral, 19 clone specific, and 155 clade specific. The topology of the breakpoint phylogeny was identical to the SNV phylogeny and counts of breakpoints and SNVs along specific branches were highly correlated (Figure 4), p-value<1.5e-6 Spearman rank).

The phylogenetic congruency of SNVs and breakpoints suggest the cell sub-sets inferred from copy number profiles represent accurate genomic clones with unambiguous genomic structure to a first approximation. While many methods have been developed for whole genome clonal deconvolution (Ha et al., 2014; McPherson et al., 2017b), most suffer from unidentifiability challenges induced by the combinatorial interaction between tumour content, cancer cell fraction, baseline ploidy and copy number genotype. Our results here demonstrate a significant step to improving clonal inference through single cell demultiplexing and clustering. We suggest this reduces the computational burden and uncertainty in inference imposed by bulk WGS methods while enhancing biological and phylogenetic interpretation of the data.

### Identification of cell type differences in whole chromosome aneuploidy and genome replication states

Bulk genome analysis of malignant and non-malignant tissues does not easily permit identification of replication states, nor the study of rare, potentially negatively selected chromosomal aberrations such as mitotic segregation errors. Mitotic mis-segregation can be observed in single cell genome sequencing as non-clonal gains and losses of whole chromosomes and we set out to examine the rate and patterns of mis-segregation across different cell types. Initial inspection of massively scaled DLP+ libraries from unsorted diploid cells (184-hTERT, GM18507) identified a minority (∼<5%) of cells with a mostly diploid genome (Figure 5a), but with aneuploidy of one or more whole chromosomes, indicating a chromosome segregation error. To quantify such events, we first clustered cells with shared copy number profiles, and then quantified outlying cells in each cluster that differ by 1 or more whole chromosome gain or loss (defined as >90% of the chromosomal length). This results in a distribution of the size and chromosomal representation of such events, over cell types (Figure 5a-g). We observed that mitotic error rates differ between different cell types, with the highest event rate of 5.2% in 184-hTERT wildtype and p53 null female cell lines (106/2038 genomes, 255/4918 genomes) whereas the reference GM18507 male cell line exhibited 2.6% events (57/2160 genomes). In contrast, the tissue derived DLP+ libraries of human follicular lymphoma and a mouse transgenic generated sarcoma model exhibited much lower rates of whole chromosome aneuploidy (6/858, 0.6% and 7/589, 1.2%, respectively), indicating that tissues and cell types vary markedly in this process. The difference between cell line and tissue derived libraries is consistent with the notion that mitotic mis-segregation rates are lower in tissues than cell lines (Knouse et al., 2014; 2018). We next asked how whole chromosome gains and losses are distributed across the genome, considering libraries where sufficient events were present to define a quantitative distribution (GM18507, 184-hTERT wildtype, 184-hTERT isogenic p53 null). We observed that (Figure 5b-g) whole chromosome gains tend to predominate over losses and are evenly distributed over the genome in both location and event sizes, for both the lymphoid cell type and breast epithelial 184-hTERT cell type. Interestingly in the isogenic p53 null breast epithelial cells, although the overall rate of whole chromosome aneuploidy is similar to isogenic wildtype (∼5.2%), the event type relationship is reversed, with losses slightly in excess of gains over all chromosomes (Figure 5e-g). The distribution of event size (number of chromosomes affected) and chromosome is similar and uniform with respect to chromosome, with the exception of chr17/19 in p53 null cells (<0.2%, Figure 5g). The latter arises from a clonally dominant Chr17 translocation which is not therefore counted as a whole chromosome in analysis.

**Figure 5.**
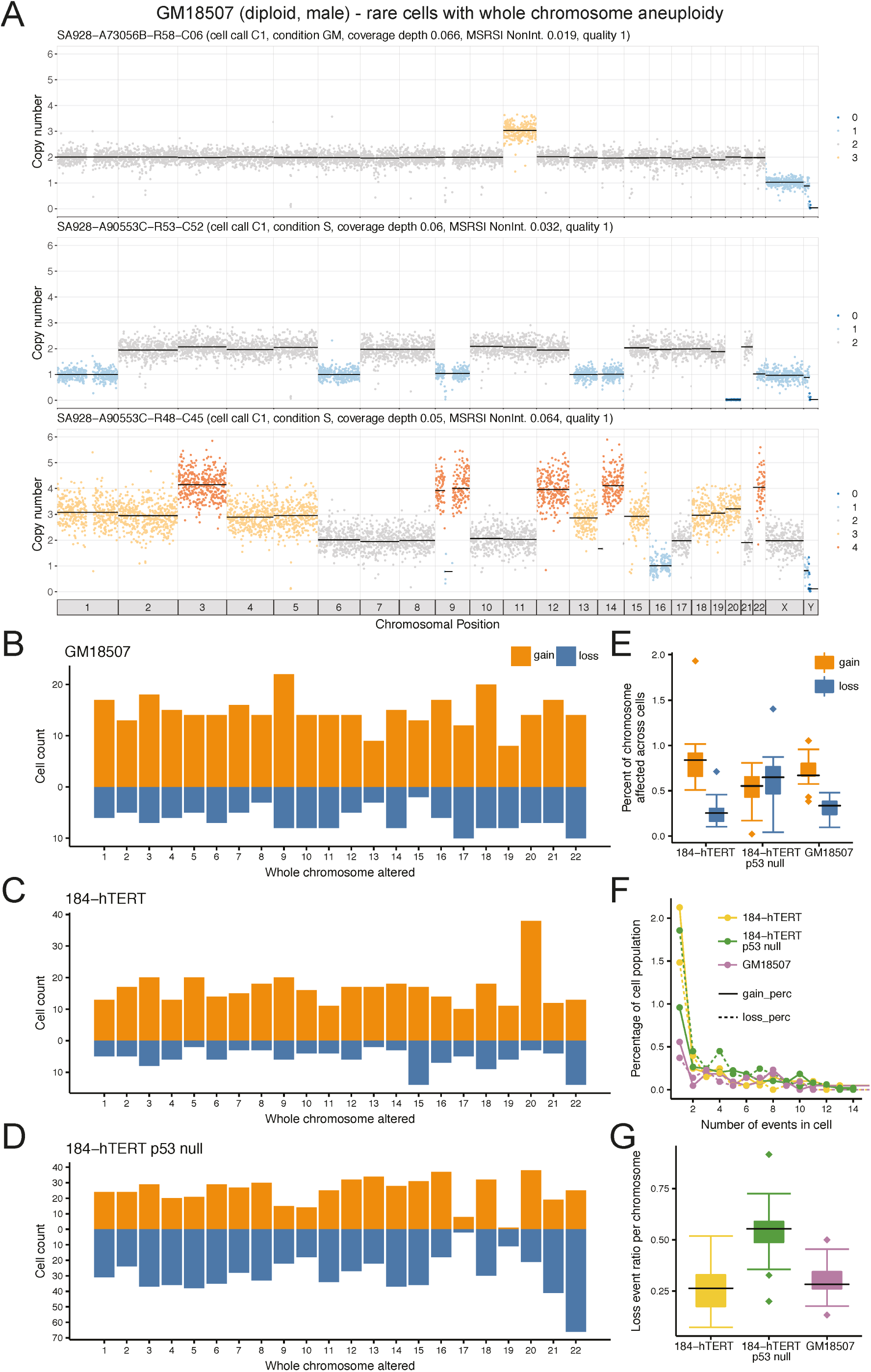
Single whole chromosome aneuploidies in single cell genomes. **a** Three examples of cells from diploid cell types exhibiting whole chromosome gain or loss patterns. b Quantification of single chromosome gain and loss patterns in diploid cell types. Left panel, vertical axis, chromosomal gains (orange) and losses (blue), horizontal axis chromosome number, in single GM18507 lymphoid cells. c As for panel c, cell type 184-hTERT. d As for panel c, cell type 184-hTERT/p53-/-. e Percentage of each chromosome affected by whole chromosome gains (orange) and losses (blue) across all cells in 184-hTERT, 184-hTERT p53 null, and GM18507. Boxplots show median and quartiles, the whiskers show the remaining distribution, dots represent outlier chromosomes. f Event number per cell (horizontal axis), for gains (solid line) and losses (dotted line), vertical axis, percentage of cells affected. Line colours represent the three cell types in the key. g Loss event ratio (losses versus gain) per chromosome for 184-hTERT, 184-hTERT p53 null, and GM18507, showing the higher rate of losses in 184-hTERT p53 null. Boxplots show median and quartiles, the whiskers show the remaining distribution, dots represent chromosomes with outlier loss ratios.

Finally we set out to address whether other genome states, such as intermediate states of replication, can be identified in tissuesfrom single-cell genome sequences in a subset of DLP+ libraries sequenced to high genome coverage per cell. Genome replication occurs asynchronously in human cells and moreover, early and late replicating regions are known to have a different GC content (Woodfine et al., 2004; Hansen et al., 2010). Partially replicated genomes are thus indicative of cells in an S-phase state, as the genome replicates asynchronously. We reasoned that variations in genome coverage and GC distribution should reflect the genome replication states. To establish the relationship, we flow sorted diploid GM18507 cells from asynchronously growing cultures, gating by DNA content and viability on cell cycle phases (Figure S12a-f). Each sorted fraction was subjected to DLP+ (G1 n = 437, S n = 393, G2, n = 359, dead n = 512) with sequencing at high depth (mean 2238604 reads per cell, ∼ 0.05x genome coverage). As expected, the distribution of GC content over binned read counts reveals a strong GC bias in S-phase cells (Figure 6a) but not in G1 or G2 cells. This distribution is also visible in the form of GC regression curves for each cell (Figure 6b). The additional mass of partially replicated DNA pushes the modal S-phase curve well above that of G1-phase cells (Figure 6a,b s-phase panel). Adequate copy number analysis of genomes requires GC correction, however the extreme GC bias in S-phase cells leads to artefactual correction (Figure 6a, second column) with standard GC normalization, due to the additional DNA content/mode of partially replicated genomes. We therefore developed a modal regression correction model (methods) which accounts for a replication-state specific GC content distribution. This results in appropriate normalization even in highly GC skewed libraries (Figure 6a, third column). Modal regression also allows for the identification of ploidy states, for example the X chromosome reads in the male GM18507 cell line, coloured purple in (Figure 6a). Once modal regression was applied, cells in S phase could be easily recognized from their partially replicated genome copy number profiles, with early replicating regions at higher copy number state than late replicating regions (Figure 6c all chromosomes, Figure 6d chromosome 4 expanded view). The pattern of representation in S-phase mirrors that of conserved early replicating regions (coloured orange Figure 6c, d)(Hansen et al., 2010) and the proportion of conserved early phase genome is much higher in S-phase cells than other states. We note that standard DNA content based flow sorting G2 phase gating is slightly imperfect and identifies some G2 state cells that are in fact still in late replication (Figure S12g). The modal ploidy of G2 states is unidentifiable in this representation, as coverage is normalized for read abundance over cells and the only hallmark of G2 states is twice the number of reads.

**Figure 6.**
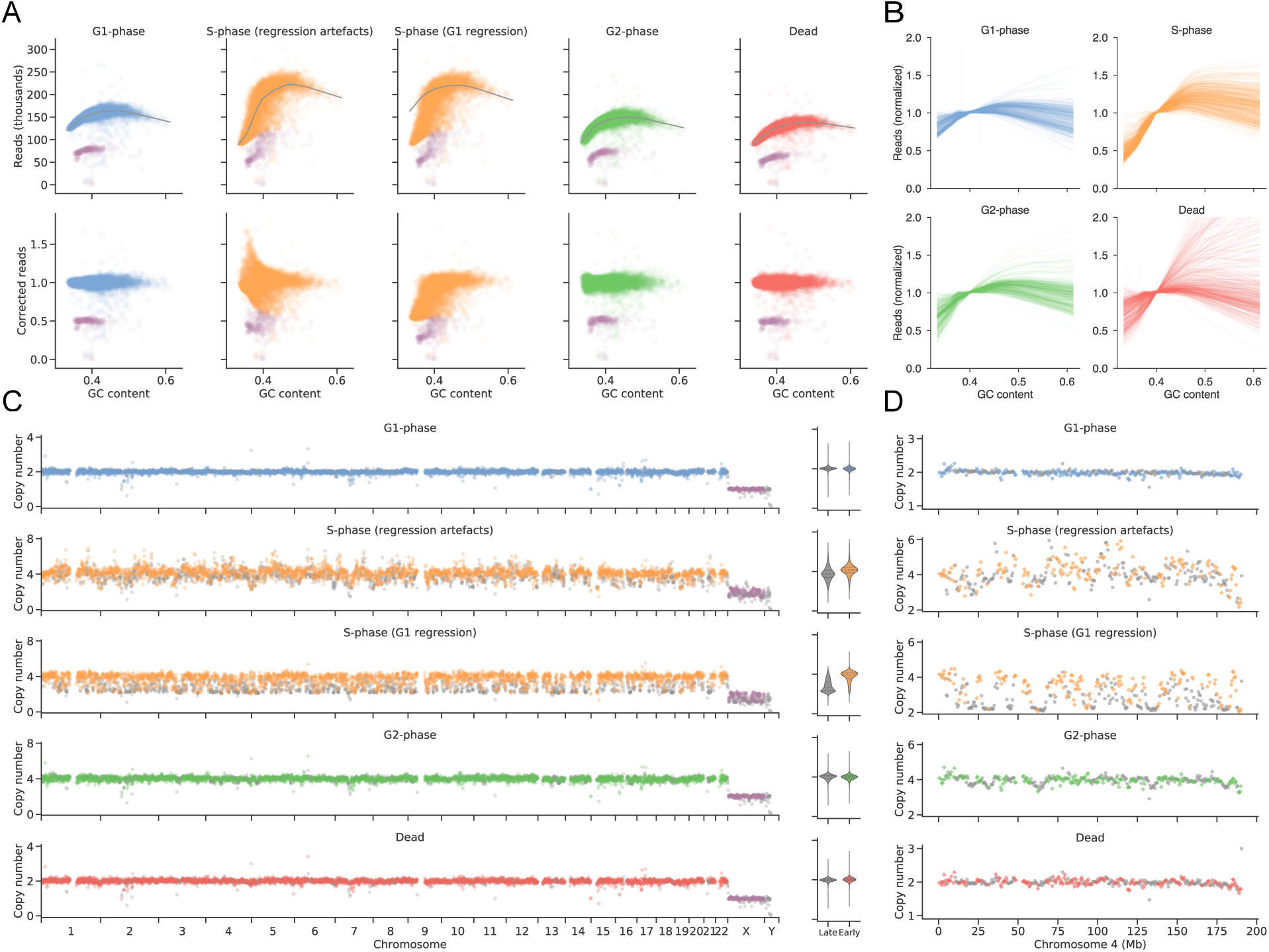
Sequencing of cell cycle sorted populations from a diploid lymphoblastoid cell line reveals early-replicating regions (n = 1701). **a** GC bias correction for merged GM18507 genomes from each flow sorted cell cycle state reveals S-phase GC bias correction artefacts. Bins from X and Y chromosomes are shown in purple. **b** Single-cell GC bias regression curves reveal S-phase cells consistently exhibit a steeper slope due to early-replicating regions with high GC content. **c** Ploidy-corrected read counts for the merged GM18507 genomes from each state (G1 n = 437, S n = 393, G2, n = 359, dead n = 512) reveal early replicating regions in S-phase. Coloured points (diamonds) denote previously characterized early replicating regions (Hansen et al., 2010), bins from X and Y chromosomes are shown in purple, while grey points (circles) denote late replicating regions. Violin plots show the distribution of late and early replicating regions for 2-copy regions. **d** Ploidy corrected read counts for chromosome 4 of the merged GM18507 genomes from each state.

To determine whether genome replication states can be effectively determined in aneuploid genomes using this approach, we also flow sorted the hypotriploid T-47D human breast cancer cell line into cell cycle fractions (G1 n = 571, S n = 625, G2 n = 807, dead n = 1039) and sequenced the genomes with DLP+. The resulting additional copy number/ploidy states over all cells are clearly visible (Figure 7a, Figure S12h) as multiple modes. Using the same modal GC regression for correction, we observed the same distribution of early and late replicating regions as in the GM18507 line, demonstrating our ability to detect S-phase in aneuploid cells (Figure 7c, d). We note that although dead cells also have a high GC bias, they are clearly distinguishable from the form of genome representation, in addition to the form of the GC bias (Figure 7b, d). Taken together, the data show that rare chromosomal aneuploidy states that do not amplify in populations and replication states can be clearly identified when single-cell genomes are sequenced at depth.

**Figure 7.**
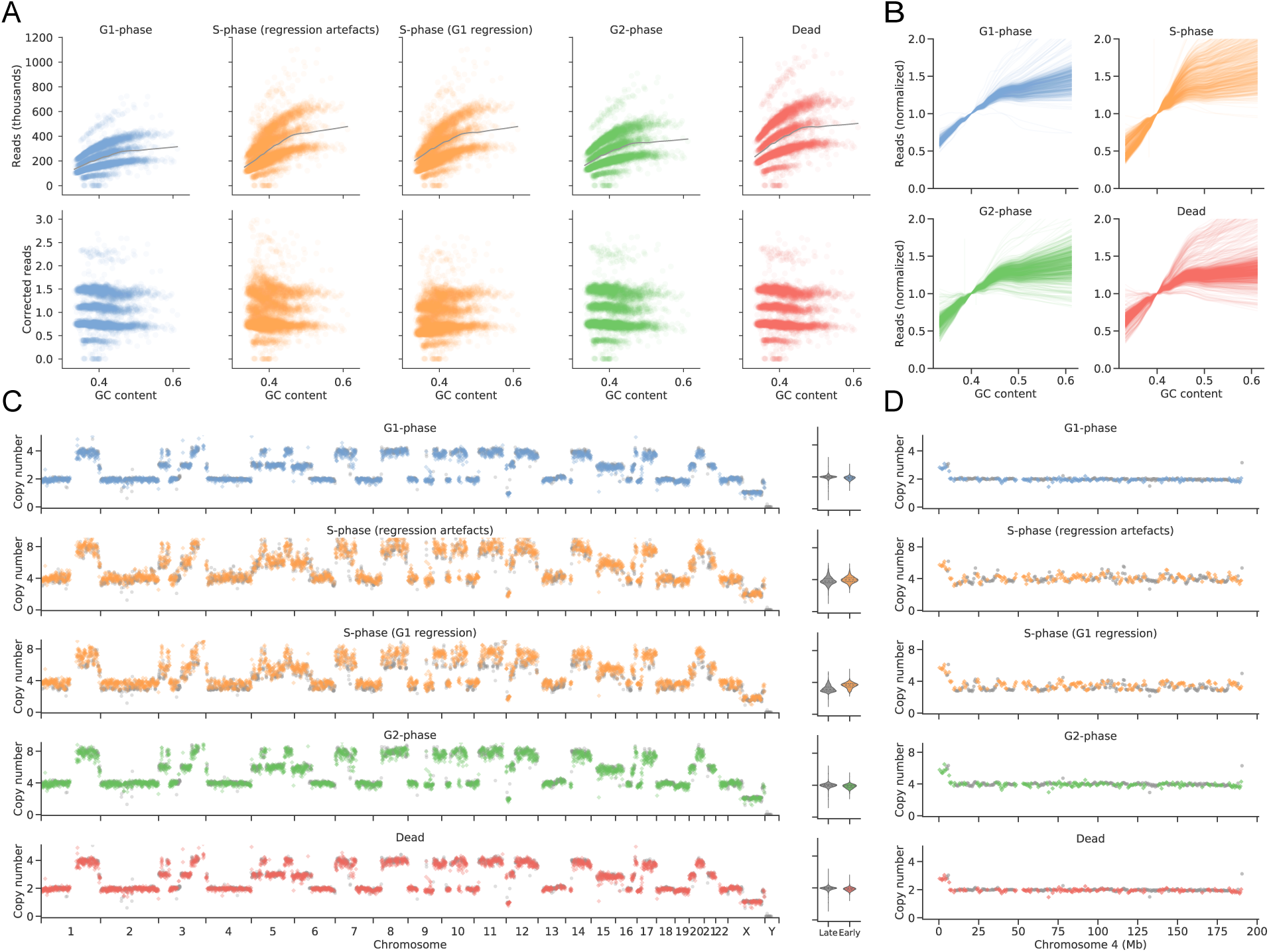
Sequencing of cell cycle sorted populations from the T-47D breast cancer cell line reveals early-replicating regions (n = 3202). **a** GC bias correction for merged T-47D genomes from each flow sorted cell cycle state reveals S-phase GC bias correction artefacts. b Single-cell GC bias regression curves reveal S-phase cells consistently exhibit a steeper slope due to early-replicating regions with high GC content. c Ploidy-corrected read counts for the merged T-47D genomes from each state (G1 n = 571, S n = 625, G2, n = 807, dead n = 1039) reveal early replicating regions in S-phase. Coloured points (diamonds) denote previously characterized early replicating regions (Hansen et al., 2010), while grey points (circles) denote late replicating regions. Violin plots show the distribution of late and early replicating regions for 2-copy regions. d Ploidy corrected read counts for chromosome 4 of the merged T-47D genomes from each state.

## Discussion

Single cell biology is opening up understanding of physiology and disease, however most of the progress and data available to date stem from single cell RNA template measurements and single cell genome analysis has lagged significantly. Scaling of single cell whole genome sequencing to tens of thousands of cells promises to accelerate the study of genome biology in normal and malignant tissues by identifying and characterizing genomic states not readily observable in bulk populations, such as rare cell populations, negatively selected background mutations and partially replicated genomes. The present resource of single cell genomes sequenced from multiple tissue and cell types illustrates that high fidelity single cell genome analysis can be conducted at scale, using commodity hardware and off the shelf reagents. Although single cell spotting can be achieved with other methods such as FACS, the small shear volumes in DLP+ minimize contamination in the carrier fluid compared to single cells isolated by FACS (piezo dispenser ∼400pL vs. FACS ∼2nL droplet volume), and the small reaction volumes substantially reduce library preparation costs compared to plate-based formats. Image information acquired during cell spotting and from whole-chip fluorescence scans can be linked to single-cell libraries or used to selectively process only cells of interest. Moreover, spotting of image identified cells or nuclei more efficiently utilizes open arrays than Poisson dilution loading (Leung et al., 2016; Gao et al., 2017; Wu et al., 2015; Goldstein et al., 2017) and greatly reduces cell doublets. To aid future reimplementation of DLP+ and deployment over a wide range of cell types, we have defined the optimal working ranges of key physical and molecular reaction parameters to obtain even genome coverage without the need for genome preamplification. It should be emphasized that a key aspect of scaling is the data processing required for interpretation and visualization of thousands of single genomes. We have implemented, from raw data, an end to end computational platform which automates calculation of quality control parameters, probabilistic classification of successful libraries, a workflow for copy number inference including GC content adjustment and an interactive, browser-based data visualization engine which allows for millisecond interaction speeds even on millions of datapoints. The cloud-based implementation of our platform facilitates virtually limitless scaling and, importantly, a data dissemination vehicle for sharing data with the broader scientific community. We anticipate other lab implementations of DLP+ will take full advantage of our software, thereby facilitating data aggregation across multiple groups.

Using DLP+, we have initiated a resource of >40,000 single cell genomes from a variety of human and mouse cell types of different cell sizes (ranging from 5 to 80 microns), malignant and non-transformed, which we have characterized by genome states. An important economic and experimental tradeoff in scWGS, given that the highest costs are still DNA sequencing reagents, is the analysis of fewer cells to greater depth of genome coverage vs shallow sequencing of many cells, borrowing strength for in depth analysis of clones identified from analysis over all cells. Small-scale events, such as SNVs and breakpoints, can be investigated at the clonal level by first identifying and merging single-cell genomes that are defined as clonal, based on shared copy number or structural events at the population level. Here we show that in aneuploid subclone containing populations, effective single nucleotide resolution can be easily achieved by merging clones defined by higher order structure such as copy number. Moreover clone specific events such as copy neutral loss of heterozygosity that cannot be easily identified in bulk populations with computational deconvolution approaches are easily identified even in minor cell populations. Thus DLP+ permits leveraging of shallow sequencing to sample thousands of cells cost effectively, rather than sequencing fewer cells at greater depth.

Using the resource presented we have also investigated two general properties of single genomes that cannot be easily obtained from bulk tissue/cell population analysis. First, we show that whole chromosome aneuploidy, which occurs at low prevalence and does not result in clonal amplification, is visible as whole chromosome gains and losses at a low prevalence in all cell types, are variable across different cell types and genotypes. Quantification of 431 such genomes out of 10963 analysed in this way across three cell lines and two tissue derived libraries is consistent with the notion of lower aneuploidy rates in tissues compared with cell lines (Knouse et al., 2014; 2018). We also observe that p53 does not appear to alter the overall event rate, consistent with the notion that whole chromosome aneuploidy may not trigger a strong p53 response (Soto et al., 2017), however, the event type is significantly altered, from chromosome gains to a slight dominance of chromosome losses. Finally, we show that partially replicated genomes can be easily identified as distinct from other biological states, or dying cells. With naive classification of single cell libraries such genomes would be filtered out, removing the possibility of identifying and characterizing such states. However we show that the distinct profiles of such genomes allows them to be identified, providing access to an important parameter of population evolution in normal and malignant cells.

In conclusion, the DLP+ scWGS platform and its derivative datasets will permit new insights into genome heterogeneity, mutational processes and clonal evolution in mammalian tissues and human disease, at scale.

## Contributions

SA, SPS, CH conceived of the research, organized the research, wrote and reviewed manuscript. EL, HZ, AS, AM-cPh, DL, developed and tested methods, performed analysis, wrote the manuscript. JBr, JBi, BW carried out tissue preparation, DLP+ library construction and sequencing. TM developed the 184-hTERT p53 null line 95.22. CN, SL, VB, MS, OG developed and deployed Montage and cellmine.org. PE, FK, TRdA, SRL performed PDX transplants and passaging. SPo developed the microscope image analysis SmartChipApp. MJT, JN, SV-W, NA, MW developed Colossus. DG, CH, TC, PW, SC developed and ran the single cell analysis pipeline. LM, RWS, TMU developed the mouse synovial sarcoma model, isolated and supplied the cells. EC and CS provided follicular lymphoma samples. RJNC and HZ developed custom parts for DLP+. DDC provided LabView assistance. SPl, YM, RC, RM, AJM, MAM assisted with sequence generations and single-cell experiments.

## Acknowledgements

The work described and the laboratories of SA and SS supported by BC Cancer Foundation, Canadian Institutes for Health Research (CIHR), Canadian Cancer Society Research Institute (CCSRI), Terry Fox Research Institute (TFRI), Canadian Foundation for Innovation (CFI), Canada Research Chairs program, Michael Smith Foundation for Health Research (MSFHR), Cancer Research UK Grand challenge IMAXT award (CRUK).

## List of Figures

**Figure 1**. Concept schematic of the experimental and computational processes for DLP+

**Figure 2**. Optimization of DLP+ scWGS library construction for the open-array format

**Figure 3.** DLP+ across different tissue types split by viability

**Figure 4.** Features from merging of clones of OV2295, OV2295(R2), TOV2295(R) cell line based on single-cell CNV

**Figure 5.** Stochastic whole chromosome aneuploides in diploid populations

**Figure 6.** Sequencing of cell cycle sorted populations from a diploid lymphoblastoid cell line reveals early-replicating regions

**Figure 7.** Sequencing of cell cycle sorted populations from the T-47D breast cancer cell line reveals early-replicating regions

## List of Supplementary Figures

**Figure S1**. Spotter setup and single-cell isolation relating to method details

**Figure S2**. Additional optimization data relating to Figure 2

**Figure S3**. DLP+ produces high quality libraries from cells and nuclei, while dead cells drop out with low read count, relating to Figure 3

**Figure S4**. Analysis work flow and classifier feature ranking relating to methods for quantification and statistical analysis

**Figure S5**. Montage single-cell data visualization relating to methods for Montage for single cell visualization

**Figure S6**. Pseudo-bulk supplementary analysis relating to Figure 4

**Figure S7**. OV2295 bulk copy number relating to Figure 4

**Figure S8**. OV2295 bulk allele ratios relating to Figure 4

**Figure S9**. OV2295 clone copy number relating to Figure 4

**Figure S10**. OV2295 clone allele ratios relating to Figure 4

**Figure S11**. Pseudo-bulk analytical workflow relating to Figure 4

**Figure S12**. Flow sort gating for cell cycle analysis of G1, S, G2 phase and dead cells by DLP+ relating to Figure 6 and Figure 7

## List of Supplementary Tables

**Table S1**. Conditions for single cell library construction

**Table S2**. DLP+ single cell sequencing metrics per single cell library

**Table S3**. Omnibus table of statistical comparisons of the DLP+ dataset

**Table S4**. Summarization of high quality cell sequencing metrics

**Table S5**. Table of SNVs per clone for OV2295 samples

**Table S6**. Table of breakpoints per clone for OV2295 samples

**Table S7**. Table of allele specific copy number per clone for OV2295 samples

**Table S8**. UMAP and GM clustering by copy number of OV2295 samples

## Methods

### Contact for reagent and resource sharing

Further information and requests for resources and reagents should be directed to and will be fulfilled by the Lead Contact, Dr. Sam Aparicio.

## Experimental model and subject details

### Cell culture

Cells from the immortalized normal human male lymphoblastoid cell line GM18507 (Coriell Cell Repositories) were cultured at 37 °C and 5% CO_2_ in RPMI-1640 medium with 2.05 mM L-glutamine (HyClone) supplemented with 10% FBS (Gibco/Invitrogen). Cells from immortalized normal human female breast epithelial cell line 184-hTERT L9 were cultured at 37 °C and 5% CO2 in MEBM Mammary Epithelial Cell Growth Medium (Lonza) with transferrin (Sigma) and isoproterenol (Sigma), supplemented with Lonza MEGM(tm) Mammary Epithelial Cell Growth Medium Singlequots. The parental 184-hTERT-L9 breast epithelial cell line, which is immortalized but not transformed and retains 3-D differentiation capacity and a diploid genome in early passages, was cultured as previously described (Burleigh et al., 2015). We generated an isogenic p53 null sister cell line using sgRNA/CRISPR targeting of the locus, which was verified by Western blotting and sequencing, resulting in the line 184-hTERT-L9-95.22. Cell line passages from the original monoclonal isolation of each cell line was recorded. Cells from the immortal human female epithelial cervical adenocarcinoma cell line HeLa (ATCC) were cultured as recommended by ATCC, at 37 °C and 5% CO_2_ in Eagle’s Minimum Essential Medium with 10% FBS. Cells from the human female high-grade serous ovarian adenocarcinoma cell lines OV2295, OV2295(R2) and TOV2295(R) (L6tourneau et al., 2012) were cultured at 37 °C and 5% CO_2_ in a 1:1 mixture of Media 199 (Sigma M5017) and Media MCDB 105 (Sigma M6395) on Corning plastics, with the media prepared as follows: Media 199 powder was dissolved in 700 mL water, stirred for 10 minutes, 2.24 g of NaHCO_3_ added, brought to 1L with water and filter sterilized. Media 105 powder was dissolved in 700 mL water, stirred 10 minutes, 14 mL of 1N sterile NaOH added, brought to 1L with water and filter sterilized. Cells from the human female breast ductal carcinoma cell line T-47D (ATCC) were cultured at 37 °C and 5% CO_2_ in RPMI-1640 Medium with 10% FBS. Cells were grown to near confluence, trypsinized, resuspended in cryopreservation media and frozen down at −1 °C/minute to −80 °C. We test the cells for mycoplasma with h-IMPACT II human pathogen testing (IDEXX Bioresearch).

### Biospecimen collection and ethical approval for patient derived breast xenografts

Tumour fragments from women diagnosed with breast lump undergoing surgery or diagnostic core biopsy were collected with informed consent, according to procedures approved by the Ethics Committees at the University of British Columbia. Patients in British Columbia were recruited and samples collected under tumour tissue repository (TTR-H06-00289) protocol that falls under UBC BCCA Research Ethics Board.

### Tissue processing for patient derived xenografts

The tumour materials were processed as previously described in (Eirew et al., 2015). Briefly, tumour fragments were minced finely with scalpels, then mechanically disaggregated for one minute using a Stomacher 80 Biomaster (Seward Limited, Worthing, UK) in 1-2 mL cold DMEM-F12 medium. Aliquots from the resulting suspension of cells and clumps were used for xenotransplantation or cryopreserved for single-cell analysis in DMEM-F12 medium with 40% FBS and 10% DMSO. Tissue was dissociated to single cells by enzymatic digestion. Cryopreserved stomached cells/organoids were thawed rapidly in a 37 °C water bath, topped up to 1.5 mL with DMEM (Sigma) and centrifuged (1100 rpm, 5 minutes), discarding the supernatant to remove DMSO from freeze media. 0.5 mL collagenase/hyaluronidase (StemCell) was added to the tissue and topped up to 1.5 mL with DMEM, pipetting up and down to dislodge tissue pellet. The tissue was incubated at 37 °C for two hours, pipetting up and down the sample every 30 minutes for 1 minute during the first hour, and every 15-20 minutes for the second hour, before centrifuging (1100 rpm, 5 minutes) and removing the supernatant. The tissue pellet was resuspended in 0.5 mL trypsin, pipetted up and down 1 minute, topped up with FBS to 1.5 mL and centrifuged (1100 rpm 5 minutes), discarding the supernatant. 1 mL dispase (StemCell) was added to the tissue pellet and pipetted up and down 1 minute, and centrifuged for 5 minutes at 1050-1100 rpm, discarding the supernatant. Digested cells were resuspended in PBS + 0.04% BSA in appropriate volume to achieve a concentration of 1 million cells/mL). Cells were passed twice through a 70 pm filter to remove remaining undigested tissue and this single-cell suspension was used for DLP+.

### Mouse model development and tissue processing

The mouse model of synovial sarcoma used herein (Martin ***et al.,*** in preparation) is based on the Haldar et al. (2007) hSS2 model which contains a conditional SS18-IRES-EGFP allele knocked into the ***Rosa26*** locus. Animals were maintained and experimental protocols were conducted in accordance with approved and ethical treatment standards of the Animal Care Committee at the University of British Columbia.

At clinical endpoint mice were humanely euthanized and the tumour was removed from surrounding tissue, and subsequently dissociated using mechanical and enzymatic digestion. To enrich for tumour cells from this mononuclear suspension, dissociated cells were stained using antibodies against various cell surface lineage markers including CD45, CD31, Ter119, F4/80, CD11b, and CD117. EGFP+ tumour cells were sorted using a BD Influx gated on eGFP expression and negative for lineage markers. Target cells were sorted into vacuum-filtered single cell (SC) collection media (DMEM containing 5% FBS) with propidium iodide. Viable target cells were subsequently further purified, and debris reduced by sorting a second time and collected into 500 pm SC collection media.

### Tissue processing for follicular lymphoma

The Research Ethics Board Number for follicular lymphoma biospecimen collection is H14-02304. Leftovers from clinical flowed samples were collected and frozen in FBS containing 10% DMSO. Cells were thawed and washed according to the steps outlined in the 10X Genomics Sample Preparation Protocol. Cells were stained with PI for viability and sorted in the BD FACSAria Fusion using a 85 pm nozzle. Sorted cells were collected in 0.5 ml of SC collection media and this single-cell suspension was used for DLP+.

## Method details

### Robot operation

All cell and reagent spotting was carried out on a contactless spotting robot (sciFLEXARRAYER S3, Scienion, Figure S1). Pulse and voltage were adjusted before every dispensing step or routine to achieve a stable droplet. Piezo Dispensing Capillary (PDC) 70 Type 1 nozzles were used for primer dispensing, PDC 70 Type 4 nozzles were used for reagent addition, and PDC 90 Type 4 nozzles or cell-qualified nozzles were used for cell dispensing. Spotter was primed daily with fresh and degassed water according to manufacturer’s recommendation. Briefly, 700mL of 18 MQ water was filtered through a 0.22 pm filter (Millipore Express Plus). The filtered water was placed in a sonicating water bath (VWR Symphony) and vacuum applied for 30 minutes using a custom adapter lid. Following the “Prime” program prompts, the bottle containing the fresh system water was then connected to the spotter. To minimize travel time during cell spotting, a custom chip holder was mounted next to the droplet camera (Figure S1a i). All other reagent additions were carried out on a temperature controlled target holder (Figure S1a vi), either at dew-point or 4 °C. If the dew-point was below 4 °C, the relative humidity was increased to 38% ± 2%, with the exception of index primers and cell dispensing where no humidity control was used. The built-in “Find Target Reference Point” function was used to adjust for placement and rotational errors. Nozzles were removed after every spot day and all system liquid lanes run dry.

### Primer spotting and wash routine

A unique combination of two dual index primers (2.1 nL each at 20 pM) were dispensed into each well of the nanowell chip (SmartChip, Seq-Ready TE MultiSample FLEX Kit, TakaraBio, 5184 nanowells arranged in a 72 x 72 well array, 110 nL each, (Figure S1a i)) in advance of cell spotting. 144 customized i7 and i5 primers (Integrated DNA Technologies) were used, where ‘NNNNNN’ was replaced with a unique hexamer barcode (Sanders et al., 2017):
i5:5’-AATGATACGGCGACCACCGAGATCTACACNNNNNNTCGTCGGCAGCGTC-3’
i7:5’-CAAGCAGAAGACGGCATACGAGATNNNNNNGTCTCGTGGGCTCGG-3’

Primers were desalted and normalized to 100 pM stock concentration in IDTE 8.0 pH. Working plates were prepared by diluting each stock primer to 20 pM in 0.1% Tween 20 in TE pH 8.0. For primer dispensing, humidity control was not used and the primers were allowed to dry down for storage at room temperature. A custom wash routine was implemented to avoid cross-contamination of index primers during spotting (Figure S2a). The wash cycle includes a series of pump and sonication steps with 2% Tween 20 and 1% SciClean solution (Scienion).

### Cell and tissue processing

#### Cell staining and sorting for cell cycle analysis

2 million cells fresh from culture suspended in 1 mL PBS were stained with Hoechst 33342 (Invitrogen), caspase 3/7 (Essen Biosciences), and propidium iodide (PI, Sigma Aldrich) for flow sorting separation of different cell phases. Hoechst 33342 requires optimization for different cell types. For the GM18507 cell line, we used 5 pg/mLwith a 30 minute incubation at 37 °C in a tissue culture incubator, in 5 pM caspase 3/7. For the T-47D line, we used 10pg/mL with a 20 minute incubation at 37 °C, in 5 pM caspase 3/7. PI was added immediately before sorting at a final concentration of 2 pg/mL and passed through a 70 pm filter.

Flow sorting was carried out at the Terry Fox Laboratory, (BC Cancer Research Centre) using a BD FACSAria III cell sorter equipped with 375 nm, 405nm, 488 nm, 561 nm and 640nm laser. Cells were sorted into media in tubes. The flow sort gating for cell cycle analysis of G1, S, G2 phase and dead cells by DLP+ is outlined in Figure S12. We gated for cells using side scatter area (SSC-A) vs forward scatter area (FSC-A) to exclude debris (black) but not dead cells (red). We next gated for single cells on this gate, using FSC width vs FSC-A to gate out doublets. We next gated for live cells on the single-cell gate using PI vs FSC to capture the live cells which are PI low. We excluded apoptotic cells on the live cell gate by gating out Caspase 3/7 high cells. On this live non-apoptotic cell gate, we gated for the cell cycle phases using DNA content of the cells measured by Hoechst 33342 staining to sort the G1, S, and G2 phases of the cell cycle individually. We also gated dead cells using the gate for single cells established in Figure S12b, but gating on the PI high, Caspase 3/7 high dead cells. Cells from different cell cycle fractions were stained and dispensed into chips as outlined in the following sections.

#### Nuclei preparation from cells

For a subset of samples (GM18507, SA501X11XB00529, SA611X3XB00821), nuclei were prepared from single cell suspensions by doubling the volume of the cells with Nuclei EZ lysis buffer (Sigma) before staining, to compare nuclei data to cell data. These were stained and Poisson spotted as described below because of the limits of the resolution of the camera.

#### Cell staining and dilution for spotting into nanowell chips

Single cell suspensions were fluorescently stained using a combination of CellTrace CFSE Cell Proliferation Kit (ThermoFisher) and LIVE/DEAD Fixable Red Dead Cell Stains (ThermoFisher), incubating for 20 minutes at 37 °C. Cells were resuspended in fresh PBS at a concentration of 220,000 cells/mL (CelleOne dispensing) or 1 million cells/mL (limiting dilution dispensing) prior to dispensing into chips with unique dual index barcodes already dispensed in each well.

### Cell spotting

Single cells or nuclei were isolated by dispensing a limiting dilution (Poisson distribution) or using active selection during cell spotting (cellenONE). For cell/nuclei isolation by limiting dilution, stained cells or nuclei were diluted to 1 million cells/mL in PBS. 1 nL of the diluted sample was dispensed into a test array to determine the isolation rate and optimize the spotting volume to achieve optimal single-cell occupancy before the remaining wells were filled using the optimized spot volume. Under optimal conditions about one-third of wells contained a single cell or nuclei; other wells were empty or contained multiple cells/nuclei. In addition, optional spotting software (cellenONE) was used to select single cells, resulting in an almost perfect single-cell isolation rate by identifying single cells inside the dispensing nozzle and depositing the desired cells selectively into reaction chambers (Figure S1). Stained cells were diluted to 220,000 cells/mL in PBS. To help avoid imaging artefacts due to reflections or external light, the robot enclosure was blacked out with opaque panels. An automated machine learning algorithm was executed after every cell uptake to set ejection and sedimentation boundaries with a mapping density threshold between 0.25 and 0.3. cellenONE allows for thresholding on size, circularity, and elongation of cells, enabling the exclusion of doublets and debris. This enables active selection of single cells as they are dispensed, overcoming the limitations of cell isolation by limiting dilution (**Figure S1 d,e**). The following advanced settings were used: min area 20, max area 250 to 1000 (depending on cell type), circularity 1.35, elongation 2.5. In addition, the LED pulse width was increased to 10 ms. Brightfield images and particle metrics from deposited cells were saved with spatial information. Isolated cells were frozen in sealed nanowell chips at −20 °C until library preparation.

#### Chip imaging and cell calling

All nanowell chips were scanned on a 10 x inverted fluorescent microscope (Nikon TI-E). Standard stages were replaced with fast travel stages to increase speed (ASI stages fitted with an ultra-course lead screw (28mm/s)). Control software was written in LabView (LabView 2015) and images were acquired on a Grasshopper3 USB camera (Point Grey Research/FLIR). A customized image analysis software (SmartChipApp in Java) was then used to confirm single cell occupancy and acquire cell state information. Intensity and area thresholding were used to select cells of choice automatically. Additional information, such a cell state (live/dead), are linked to each well after imaging the entire device and automatically extracting fluorescent imaging information (Figure S1 f, g). Automated calling was then reviewed, and a spotting robot input file was created to process selected wells only. Since imaging occurs before the library preparation reagents are spotted, doublets, empty wells, or cells with contamination can be excluded from library preparation (Figure 1a). All imaging information together with additional information, such as sample type, sample processing, and spatial information were recorded in a custom database, Colossus.

### Library preparation protocol optimization

We used a one-pot transposase chemistry (Nextera DNA Library Preparation Kit, Illumina) as described by (Zahn et al., 2017). Following all reagent additions, nanowell chips were sealed (Microseal film A, BioRad; pressed on with a pneumatic sealer) and reagents collected at the bottom of the well with a centrifugation step at 3214 g for 2 min. All chip incubations, with the exception of the cell heat lysis, were carried out on a flatbed thermal cycler (DNA Engine Tetrad 2, MJ Research), followed by a centrifugation step for 2 min at 3214 g. Table 1.1 summarizes all experimental conditions; a detailed description of each condition can be found below.

### Dispensing method

Cells were dispensed by a limiting dilution (Poisson) to isolate single cells or single cells were selected directly in the nozzle (block cellenONE, see section Cell spotting). Active selection of cells in the nozzle results in a block pattern vs. the scattered pattern of single isolated cells resulting from the limiting dilution. We investigated the effect of sample distribution on the chip and mimicked a limiting dilution-like scattered distribution using target dispensing of selected single cells (scattered cellenONE).

### Lysis

We investigated the following lysis conditions on the open-array platform. <u>Lysis buffer ‖ Protease:</u> G2 lysis buffer was prepared with 25 pL lysis buffer G2 (Qiagen) and 2.5 pL (+) Qiagen Protease (Protease was re-suspended in 7 mL UltraPure water). Direct lysis buffer was prepared by combining DirectPCR Cell Lysis Reagent (Viagen) (25 pL), Qiagen Protease (+: 2 pL, ++: 5 pL, +++: 10 pL) in 5% glycerol and 0.1% pluronic in PCR water. <u>Volume& presoak time:</u> 1 nL or 10 nL of the specified lysis solution was dispensed into the selected wells of the nanowell chip and cells were incubated at 4 °C (0 hours, 2 hours, 4 hours or overnight (19-22 hours)). <u>Protease top-up:</u> If applicable, 2.5 nL of additional lysis solution was added to each well. <u>Water bath/temp/dry down:</u> Heat lysis was carried out at 50 °C for 1 hour followed by a protease inactivation incubation at 70 °C for 15 minutes, with a final cooling to 10 °C. If applicable, cell heat lysis was performed by immersing the sealed chip into a water bath at 50 °C for 1 hour, followed by a transfer to a thermal cycler for protease inactivation (70 °C for 15 minutes, 10 °C forever). During immersion, the chip was mounted in a custom-built chip clamp to ensure a secure fit of the seal. Finally, a dry down might have been performed at room temperature for 15 minutes, followed by dispensing 10 nL of water to equalize volumes before tagmentation. Lysis solution was added to all wells, including gDNA, no-template (NTC) and no-cell (NCC) controls.

### Tagmentation

After cell lysis, 18 nL of the 2.2 nL tagmentation mix (9 nL TD Buffer, 2.2 nL TDE1, 0.165 nL 10% Tween-20), 3.5 nL tagmentation mix (14.335 nL TD Buffer, 3.5nL TDE1, and 0.165 nL 10% Tween-20) or 6.5 nL tagmentation mix (11.3 nL TD Buffer, 6.5 nL TDE1, and 0.165 nL 10% Tween-20) in PCR water were dispensed into each well and incubated at 55 °C for 10 minutes followed by cooling to 10 °C.

### Neutralization

The tagmentation reaction was neutralized with 4 nL Qiagen Protease and 4 nL 0.2% Tween-20, and an incubation at 50 °C for 15 min, followed by a protease inactivation incubation for 15 minutes at 70 °C, with a final cooling to 10 °C.

### PCR

After neutralization, 39 nL of PCR master mix (19.5 nL NPM, 6.5 nL PPC, 0.65 nL 10% Tween-20, 12.35 nL PCR water) was dispensed to each well. PCR was performed using the following conditions: 72 °C for 3 minutes; 95 °C for 30 seconds; 8 cycles or 11 cycles of 95 °C for 10 seconds, 55 °C for 30 seconds and 72 °C for 30 seconds; 72 °C for 3 minutes; and finally 10 °C. The indexed single-cell libraries were then recovered by centrifugation through a recovery funnel into a pool. Finally, size selection was performed using a 1.8 x Ampure XP (Beckman Coulter) bead to sample ratio.

### Optimized DLP+ method

<u>Dispensing method</u> Single cells were selected directly in the nozzle using cellenONE software. <u>Lysis buffer ‖ Protease:</u> Direct lysis buffer was prepared by combining DirectPCR Cell Lysis Reagent (Viagen) (25 pL), Qiagen Protease (2 pL in 5% glycerol and 0.1% pluronic in PCR water. <u>Volume ‖ presoak time:</u> 10nL of this lysis solution was dispensed into the selected wells of the nanowell chip and cells were incubated at 4 °C overnight (19-22 hours). <u>Heat lysis:</u> Cell heat lysis was performed by immersing the chip into a water bath at 50 °C for 1 hour, followed by a transfer to a thermal cycler for protease inactivation (70 °C for 15 minutes, 10 °C forever). During immersion, the chip was mounted in a custom-built chip clamp to ensure a secure fit of the seal. Lysis solution was added to all wells, including gDNA, no-template (NTC) and no-cell (NCC) controls. <u>Tagmentation</u> After cell lysis, 18 nL of the 3.5 nL tagmentation mix (14.335 nL TD Buffer, 3.5 nL TDE1, and 0.165 nL 10% Tween-20) in PCR water were dispensed into each well and incubated at 55 °C for 10 minutes followed by cooling to 10 °C. <u>Neutralization</u> The tagmentation reaction was neutralized with 4nL Qiagen Protease and 4nL 0.2% Tween-20, and an incubation at 50 °C for 15 minutes, followed by a protease inactivation incubation for 15 minutes at 70 °C, with a final cooling to 10 °C. <u>PCR</u> After neutralization, 39 nL of PCR master mix (19.5 nL NPM, 6.5 nL PPC, 0.65 nL 10% Tween-20, 12.35 nL PCR water) was dispensed to each well. PCR was performed using the following conditions: 72 °C for 3 minutes; 95 °C for 30 seconds; 8 cycles of 95 °C for 10 seconds, 55 °C for 30 seconds and 72 °C for 30 seconds; 72 °C for 3 minutes; and finally 10 °C. The indexed single-cell libraries were then recovered by centrifugation through a recovery funnel into a pool. Finally, size selection was performed using a 1.8 x Ampure XP (Beckman Coulter) bead to sample ratio.

### Quality control and sequencing

Cleaned up pooled single-cell libraries were analyzed using the Aglient Bioanalyzer 2100 HS kit. Libraries were sequenced at UBC Biomedical Research Centre (BRC) in Vancouver, British Columbia on the Illumina NextSeq 550 (mid- or high-output, paired-end 150-bp reads), or at the GSC on Illumina HiSeq2500 (paired-end 125-bp reads) and Illumina HiSeqX (paired-end 150-bp reads).

## Quantification and statistical analysis

### Data analysis workflow

We obtained dual-index demultiplexed FASTQ reads from HiSeq instruments, and we used these FASTQ reads as input into our workflow automation pipeline. Our workflow was written in Pypeliner (https://anaconda.org/dranew/pypeliner), is publicly available (https://svn.bcgsc.ca/bitbucket/projects/SC/repos/single_cell_pipeline/browse), and outlined below. All tools were run on default settings unless otherwise specified, exact version and environment are encoded in the public available workflow link above. Workflow is outlined in Figure S4a).

#### Single-cell alignment

Reads were trimmed with `TrimGalore` to remove adapters and paired-end single-cell FASTQs were aligned with `bwa`, `aln` mode. PCR duplicates were marked using `picard MarkDuplicates` and read alignment metrics are computed for each cell.

#### Single-cell CNV calling

Reads were tabulated for non-overlapping 500k genomic regions or “bins”. A modal regression normalization was performed to reduce GC bias. Copy number was called using `HMMcopy`, under 6 possible ploidy settings and a fit was computed for each ploidy with the best fit returned per cell.

#### Data analysis infrastructure

A major challenge to analyzing the data was dealing with the number of files present. With an average of 1000 cells per library and up to 3 libraries a week being produced, new infrastructure was required to be built specifically to keep up with the flow of data.

#### Modal regression for GC bias correction

In previous studies, GC bias correction was conducted using fitted local regression curves (Zahn et al., 2017; Ha et al., 2012). When applied to samples whose average ploidy fell between integer values, this approach caused normalized read count bands to fall between integer lines, complicating ploidy estimation and integer copy number inference. To address this, a modal regression curve fitting procedure was applied as follows. For each cell, second-order polynomial quantile regression curves were computed for each quantile from the 10th to 90th using the `statsmodels` Python package. Next, the area between quantile regression curves in the 10th to 90th GC quantiles was computed by integration. A lowess local regression curve was fitted to the curve distances, and the minimum of the smoothed curve was selected as the modal quantile regression curve. To correct for GC bias, read counts for each bin were divided by the predicted value at the modal quantile curve. Corrected read counts were passed as input to `HMMcopy` for segmentation and copy number state inference.

#### Copy number calling

Copy number calling was done using `HMMcopy` as described in Ha et al. (2012). Briefly, we obtained a histogram of aligned read start positions at 500k bins resolution across the genome, corrected for the bias effect of GC content on sequencing depth, then predicted the copy number with a 12 state Hidden Markov Model. In order to optimize `HMMcopy` which was originally designed for heterogeneous bulk tumour genomes for single cells, two major changes were made. Firstly, instead of applying the default loess regression method from `HMMcopy`, we implemented a modal regression algorithm that correctly normalizes bin counts to integer values as expected of single-cell profiles. Secondly, instead of a 7 state model, we expanded to a 12 state model to better capture the dynamic range of copies we encounter in single-cell libraries.

Determining the correct copy number calls was complicated by the lack of a “ladder” to map observed coverage depth to biological chromosome count. This means that the data from a perfectly normal female human diploid cell could also be explained by a single haploid genome, or any other ploidy genome sampled uniformly. For normal cells like this, we resolved this issue with prior knowledge and manually set the parameter set to assume diploid biological input. For cells with more events in their copy number profile, we can use these events to infer the actual copy number, under the assumption that all events should be derived from an integer number of chromosomes as data was derived from a single genome. Algorithmically, we made copy number predictions with `HMMcopy` using 6 possible ploidy assumptions (haploid to hexaploid), by multiplying the normalized data by 1 to 6. We then computed a penalty score we call *halfiness* that penalizes non-integer copy number predictions and select the ploidy that minimizes this penalty for downstream analysis.

Formally, halfiness is a single score computed for each cell independently as follows:

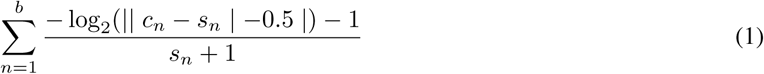

Where *b* is the number of bins in the genome, n is the median predicted copy numbers for the segment the bin resides in (where a “segment” is simply contiguous bins with the same copy number state), and s is the integer copy number state that is one of 0 to 11. Intuitively, the closer the predicted copy number is halfway between two integer copy number states, the higher the penalty score. We also penalize these errors much more heavily at lower states, as practically these are the states with the highest occurrence and confidence. We cap (|*c*_*n*_ *- s*_*n*_|) at 0.499 to prevent the asymptotic numerator from going to infinity and handle edge cases where this difference exceeds 0.5.

#### Computing cell quality

Using the `randomForest` package from CRAN (https://cran.r-project.org/web/packages/randomForest/index.html), we created a binary classifier trained on 20000 labelled cells. The labelling was done by two of the co-authors independently on separate sets of cells (13000 and 7000). A total of 6700 were labelled as good and 13300 as bad, based on the integerness of the segments and overall noise in the single cell-library, with roughly 2500 other cells discarded during the process due to ambiguity. This classifier produces a quality score (QS) for each cell using 18 cell specific features ordered by importance (Figure S4b).

- `total_mapped_reads:` the total number of mapped reads
- `total_duplicate_reads:` the total number of duplicate reads
- `MBRSM_dispersion:` median of bin residuals from segment median copy number values
- `MSRSI_non_integerness:` median of segment residuals from segment integer copy number states
- `scaled_halfiness:` a scaled metric to assess integer goodness of fit, as previously defined
- `MBRSI_dispersion_non_integerness:` median of bin residuals from segment integer copy number states
- `breakpoints:` number of intrachromosomal breakpoints
- `loglikehood:` log-likelihood of HMMcopy CNV fit
- `mad_hmmcopy:` mean absolute deviation of CNV results
- `total_halfiness:` halfiness, but not scaled by copy number state (no denominator in definition)
- `cv_hmmcopy:` coefficient of variation of CNV results
- `mean_state_mads:` the mean across all MADs of each copy number state
- `mean_state_vars:` the mean across all variances of each copy number state
- `percent_duplicate_reads:` percentage of reads that are duplicates
- `autocorrelation_hmmcopy:` autocorrelation of CNV results
- `mean_copy:` mean copy number of all bins
- `standard_deviation_insert_size:` read insert size standard deviation
- `state_mode:` the most commonly occurring copy number state

The random forest tree was trained using all default settings, where 500 trees were grown with 4 variables tried at each split, and a training out-of-bag estimate error rate of 2.38%. The quality score is the probability of being in the “good” class, and ranges from 0 to 1, with 1 indicating a high probability that a cell is high quality.

The strongest two features were representations of sequenced depth, with the next 4 all being various calculations of how well the copy number profile fits to integer copy number states. On the other end, it is somewhat reassuring to the generalizability of the model that the actual mean and mode copy numbers are not significant features in determining cell quality.

When applied to the entire dataset the resulting quality distribution is strongly bimodal, with 34% of results are below 0.1 and 52% above 0.9. Given this distribution, as visualized in Figure S2a, we used a threshold score of ≥ 0.75 to capture a highly confident subset of cells for downstream analysis.

### Montage for single cell visualization

Data from the single-cell analysis pipeline was loaded into Montage for visualization and interpretation. Montage is a JavaScript web interface, built using D3.js. Driven by an Elasticsearch back-end and deployed as an Elasticsearch plugin, Montage enables responsive querying of millions of data points. Detailed documentation and the complete source code are available on GitHub - https://github.com/shahcompbio/montage.

### Pseudo-bulk analysis

We sought to use DLP as a platform for reconstructing the clonal architecture and evolutionary histories of sequenced samples. Copy number changes are highly homoplastic, potentially confounding phylogenetic reconstruction. SNVs and breakpoints are more optimal phylogenetic markers as their low homoplasy allows application of the infinite sites hypothesis. However, the low per-cell coverage obtained with DLP is insufficient for either calling or assessing presence of SNVs or breakpoints in single cells. We thus sought to reconstruct clonal architectures using a two step procedure, first clustering cells by their copy number profiles, then treating the resulting clusters as pseudo-bulk genomes in a multi-sample analysis of SNVs, breakpoints, and loss of heterozygosity.

#### Clustering by copy number

Accurate clustering is a crucial first step of pseudo-bulk analysis. If cells from divergent phylogenetic clades are clustered together, downstream analysis will be unable to accurately assign SNVs to the correct clone. In the extreme case, a poor clustering will introduce a fraction of contaminating cells in each cluster, resulting in SNVs being called as present in all clones and obscuring any phylogenetic signal. We thus chose a method of copy number clustering with stringent filtering of outliers and low quality clones.

We first used UMAP version 0.2.3 (McInnes and Healy, 2018) with default parameters to produce a reduced 2 dimensional representation of the copy number data. We then clustered the resulting reduced dimensionality data using a Gaussian Mixture Model, over-specifying the number of clusters at 20. Clusters composed of less than 50 cells were excluded from further analysis. Cells with an RMS of 0.8 from the median copy number of the cluster were excluded from further analysis. The median copy number of the cluster was used as the measurement of total copy number for each cluster.

#### Allele specific copy number

We computed allele specific copy number using a previously described approach (McPherson et al., 2017b), detailed below. In a matched normal sample we measured reference and alternate allele counts for SNPs from the thousand genomes phase 2 reference panel. We used a binomial exact test to filter for SNPs heterozygous in the normal sample. Using shapeit (Delaneau et al., 2011) and the thousand genomes phase 2 reference panel, we computed haplotype blocks.

Next we measured per cell reference and alternate allele counts for heterozygous SNPs in the DLP data. Per clone counts were aggregated by summing across cells in each cluster. Haplotype blocks that were split at boundaries of `HMMCopy` bins, and major and minor haplotype allele counts were computed for each cluster and each haplotype block. We then used an HMM with Binomial emission to model haplotype block counts, and used the viterbi algorithm to compute the optimal minor copy number state per bin. To account for outliers, the emission was a mixture of a Binomial and a uniform distribution. Specifically, given observed total copy number *t*, unobserved minor copy number *z*, minor haplotype allele counts *x* and total haplotype block counts *k*, the likelihood is given by Equation 3. We used a fixed transition matrix favouring self transitions as given by Equation 4, where s is the maximum copy number state.

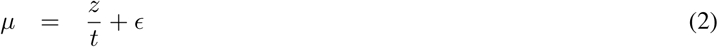

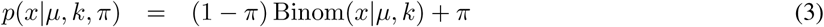

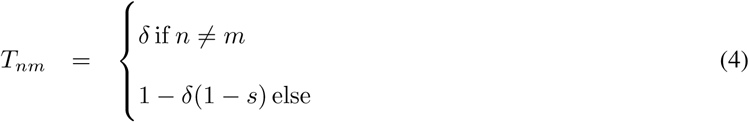

For the purposes of this study we fixed the parameters at *ε* = 0.01, *π* = 0.01, *δ* = 1e - 4 and *s* = 11.

#### SNV and breakpoint calling

We used mutationseq (Ding et al., 2012) and strelka (Saunders et al., 2012) to call SNVs in merged DLP genomes. We first created merged BAMs for each DLP library, split into non-overlapping 10MB regions. Both mutationseq and strelka were used to call SNVs with default parameters as for SNV calling in bulk whole genome data, as previously described (McPherson et al., 2016). We generated a set of high quality SNV predictions by filtering for SNVs with strelka somatic score ≥ 20, mutationseq probability ≥ 0.9, envode 50mer mappability ≥ 0.99. Samtools was used to extract per cell, per SNV reference and alternate allele counts for a union set of filtered SNVs.

For breakpoint prediction we used deStruct (McPherson et al., 2017a), which produced per cell per breakpoint counts. Breakpoints were filtered for predictions with at least 5 split reads, and at least 250 nucleotides anchoring the predicted sequence on either side of the breakpoint (template_length_min feature).

#### Phylogenetic Analysis

We used the stochastic Dollo evolutionary model in conjunction with a binomial read count likelihood to reconstruct the evolutionary relationships between clones as previously described (McPherson et al., 2016). In brief, alternate SNV counts are modelled as binomial distributed given total counts at the SNV loci. To calculate a probabilistic score for a given tree and SNV, we calculated the likelihood of the SNV reference and alternate counts, marginalizing the possible origins of that SNV throughout the tree, in addition to any SNV losses due to copy number change. Losses occur with a branch specific probability that is learned as part of model fitting. Exhaustive search is used to select the maximum likelihood tree. Given the ML tree, the origin branch and branch specific losses are calculated for each SNV by maximum likelihood.

We generated a breakpoint phylogeny using hierarchical clustering of binary presence/absence data with average linkage and euclidean distance.

## Data and software availability

### Data

The single-cell FASTQs have been deposited in the European Genome-phenome Archive under ID code EGAS00001003190.

### Software and code

We developed a suite of tools to facilitate large scale processing of DLP+ sequenced libraries on a local high performance computing cluster with the ability to burst compute with Microsoft Azure’s Batch Compute. The suite of tools includes 2 databases, Colossus and Tantalus, an application for analyzing the aluminum SmartChip, and an analytical pipeline. Colossus acts as a lab notebook for the molecular biologists, cataloguing samples, DLP+ libraries, lanes of sequencing of those libraries and per cell metadata. Tantalus, by contrast, is a system used primarily by analysts for tracking metadata of sequencing datasets, analyses and results. Sisyphus communicates with the RESTful APIs of Colossus and Tantalus to prepare inputs for analyses and execute those analyses.

#### SmartChipApp

The SmartChipApp is an interactive application that analyses captured images of cells spotted in a grid in a nanowell SmartChip. Images from two fluorescence channels are captured to highlight the state of the cells in each spotted well. For example, one channel could be used to highlight the cells that are live and the second channel could be used to highlight the cells that are dead. The application automatically detects and quantifies the number of live and dead cells in each well and saves the results in an Excel table. Cell calls can be manually revised by the user. The application also saves files that control the spotting robot, allowing wells to be selectively addressed based on their contents.

The code for the SmartChipApp is publicly available and accessible at https://github.com/shahcompbio/smartchipapp.

#### Colossus

Colossus catalogues samples, DLP+ libraries, lanes of sequencing of those libraries and per cell metadata. The implementation of Colossus uses Django web framework with a PostgreSQL database. Data can be browsed in an intuitive front end that includes search features, tabular presentations of the data, and the ability to add, edit and delete samples, libraries, and sequencings. Per cell metadata can be imported into Colossus from Microsoft Excel spreadsheets generated by the SmartChipApp. Additionally, Colossus provides the ability to generate tables required for submitting a library for sequencing and demultiplexing the sequenced library into per cell fastqs. A RESTful API allows for integration into automation scripts.

The code for Colossus is publicly available and accessible at https://github.com/shahcompbio/colossus.

#### Tantalus

Tantalus is an organizational tool for tracking DLP+ sequencing datasets and analyses. The implementation of Tantalus uses Django web framework with a PostgreSQL database. Metadata of single cell datasets, including file paths and sample and library information, are browsable and searchable in an html front end. A python celery based backend allows for the automation of tasks including data import and file transfers, with automation of analyses planned in future versions. A RESTful API allows for integration into automation scripts.

The code for Tantalus is publicly available and accessible at https://github.com/shahcompbio/tantalus.

#### Single Cell Pipeline

The analytical pipelines for processing the raw sequence data are packaged as a single python module, single_cell_pipeline. The pipelines use the pypeliner workflow orchestration tool to define dependencies between tasks and provide the ability to run the pipelines in parallel environments including multi-processing, grid engine, and in Microsoft Azure Batch. In brief, an alignment and QC pipeline generates aligned BAMs from Fastq files and runs HMMCopy for each cell. A series of additional pipelines for variant calling, germline calling, and breakpoint calling are used for pseudo-bulk analysis. Each pipeline takes as input a list of input BAMs or Fastq files per cell in YAML format and outputs a set of results tables in HDF5 format. Details of how to run the pipelines are shown in the readme available in the repo.

The code for the single cell pipeline is publicly available and accessible at https://github.com/shahcompbio/single_cell_pipeline. The code for pypeliner is publicly available and accessible at https://github.com/shahcompbio/pypeliner.

#### Cell cycle analysis

Code for cell cycle analysis is publicly available and accessible at https://svn.bcgsc.ca/bitbucket/projects/SC/repos/dlpplus_protocols/browse/cell_cycle_plots

#### Montage QC dashboard

We constructed a Montage dashboard configuration for quality control (QC dashboard) and library assessment, which consists of the following for all cells in a given analysis (Figure S5): (1) a heatmap of copy number states across the genome per cell (Figure S5c), (2) a table of experimental conditions to allow for interpretation of experiment and controls (Figure S5d), (3) a chip heatmap revealing a user selected sequencing metric (ex. the total mapped reads) per cell across the physical device (Figure S5e), (4) an interactive scatterplot of the cells’ quality metrics (which can be selected, Figure S5f) and (5) a violin plot of the distributions of metrics for each combination of cell call and experimental condition (Figure S5g). All of the plots can be filtered by clicking the green Data Filter circle (Figure S5a) and entering filters in the menu displayed. The metrics displayed in the chip heatmap, scatter plot, and violin plot can be changed by clicking the plot and using the same side menu. Code and documentation for Montage is publicly available and accessible at https://github.com/shahcompbio/montage

## Additional resources

Cellmine for single cell data visualization: https://www.cellmine.org

A guest account has been created for editors and reviewers to view cellmine.org prior to it being publicly released:

User: guest

Password: fF!uUVG6z

DLP+ protocol download: https://svn.bcgsc.ca/bitbucket/projects/SC/repos/dlpplus,protocols/browse)

## Supplemental Information

### Cost of DLP+ consumables

DLP+ is a flexible platform, a researcher can choose to target hundreds or thousands of cells in a single experiment. For the purposes of this cost analysis we will assume 1000 cells are included per library, 3000 cells per open array chip, and 6000 cells (two chips) per library construction and library quality control check.

| Component | CAD for a unit | Quantity/unit | Amount/use | Cells/use | CAD/cell |
| --- | --- | --- | --- | --- | --- |
| Primers | 1353.60 | 504 Smartchips filled | 1 chip | 3000 | 0.00090 |
| SmartChip | 270.00 | 1 Smartchip | 1 chip | 3000 | 0.09000 |
| Microseal A | 189.00 | 200 seals | 6 seals | 3000 | 0.00189 |
| CFSE | 232.50 | 180 ul | 0.4 ul | 50000 | 0.00001 |
| Live Dead Red | 415.23 | 250 ul | 0.4 ul | 50000 | 0.00001 |
| Lysis Buffer | 134.93 | 50 ml | 42.5 ul | 3775 | 0.00003 |
| Protease | 95.00 | 7 ml | 25 ul | 3775 | 0.00009 |
| Nextera Kit | 10455.80 | 1.55 ml NPM | 75 ul | 3375 | 0.14990 |
| Ampure | 1555.72 | 60 ml | 135 ul | 1000 | 0.00350 |
| Agilent HS | 849.00 | 10 Agilent HS chips | 1 chip | 6000 | 0.01415 |
| TOTAL |  |  |  |  | 0.26048 |

### Montage QC dashboard use case

The interactive data filtering and exploration features of Montage proved to be valuable for data quality assessment.

#### Montage use case examples visualize mouse cell contamination and mixed ploidy samples

Identification of contaminating mouse cells in xenograft samples: Dataset SC-899

The dimensions of the scatter plot were set to display total reads (x-axis) and total mapped reads (y-axis). The off-diagonal points quickly illustrate which cells have unusually low numbers of mapped reads, and selecting these cells reveals that they are the source of noisy copy number profiles in the copy number heatmap. In cases where PDX tissue are analyzed, this can quickly filter contaminating mouse cells from the heatmap.

## Identification of physical parameter ranges for implementation of DLP+

First, for assessment of per cell and library construction performance, we developed a series of 46 pre and post alignment sequencing metrics as features of each single cell genome. In addition to standard sequencing and alignment metrics, we developed several new quality metrics based on “integer-ness” scores of copy number states that were shown to substantially improve discrimination (Methods). We explored the weights of these features using a random forest applied to manually classified libraries from the diploid lymphoblastoid cell line GM18507 (Figure S4b). The total high quality aligned sequence reads (total mapped reads), the median of bin residuals from segment integer copy number states (MBRSI dispersion non-integerness), and the median of segment residuals from segment integer copy number states (MSRSI non-integerness), a read depth independent metric of non-integer copy number assignment, were the metrics with the highest weights in identifying poorly performing libraries. To simplify the evaluation of the library quality from hundreds of cells, we implemented a random forest classifier to jointly evaluate 18 of these sequencing and post-alignment metrics and provide a single quality score for each cell (QS; Figure S3a, sTable 1).

For initial reaction parameter exploration we built sequencing libraries from the GM18507 lymphoblastoid cell line (The International HapMap Consortium, 2005), used an HMM (Ha et al., 2012) to infer copy-number profiles, and classified cells in each library using our trained random forest classifer. We included all wells containing single cells (as identified by microscopy) in the following analysis. Cells that did not produce any reads were assigned a quality score of zero (QS = 0) during classification.

We examined limiting dilution cell dispensing against real-time selected cells (cellenONE, Scienion) dispensed in a block or limiting dilution-like scatter pattern (Figure S2d), and found no significant improvement in library quality of the actively selected cells over the passively dispensed cells for the high Tn5 concentration (KW test, p value = 0.678 (2.2 nL tn5 condition)). The overall library quality built from single cells remained poor, while libraries built from gDNA or crude lysate produced high-quality copy number profiles (data not shown), motivating further optimization. The cellenONE block spotted cells did show a significant increase in total mapped reads compared to the scattered pattern cells that were cellenONE or Poisson spotted (mean total mapped reads cellenONE block = 592240, cellenONE scatter = 207090, limiting dilution = 247197 (2.2 nL tn5 condition), KW test scatter vs block p value = 3.94e-08). This motivated a switch to cellenONE spotting for all future cell spotting.

Lysis volume, buffer type and crucially, time, proved to be one of the most important parameter sets determining overall performance. We tested new lysis buffer conditions (Figure 2c), which are a trade-off between reducing off-reactions in low volumes, vs. evaporation losses and the need to dilute buffer components in subsequent one-pot reactions. The low volumes of lysis buffer (1 nL Buffer G2, Qiagen) used in the microfluidic device (Zahn et al., 2017) proved not to be robust in the open-array format (Figure 2a, c ii 1 nL G2) because the droplets do not fully cover the well. However we found that higher volumes of the same buffer poisoned the downstream reactions due to insufficient dilution (Figure 2c ii 10 nL G2; mean total number of reads 1nL G2 buffer = 1443217, 10 nL G2 buffer = 4136). To address this, we evaluated a PCR compatible lysis buffer (DirectPCR Cell Lysis Reagent, Viagen). The new lysis buffer in combination with the increased lysis volume (10 nL; Figure 2c ii) 10 nL Viagen) significantly improved the mean library quality compared to the initial microfluidic lysis condition (KW test, p = 3.00 e-03), but the overall quality remained poor compared to the MF-DLP dataset (Figure 2b i). We next tested the PCR compatible lysis buffer with and without the addition of protease (Protease, Qiagen; Figure 2c iii, SUPPLEMENT FIG, respectively) against a low (2.2nL), medium (3.5nL), and high (6.5nL) Tn5 concentration. The overall quality score remained poor suggesting that the genomic DNA was still not fully accessible due to incomplete lysis, however, in the presence of protease in the lysis mix the library quality significantly improved for the medium (KW test, p = 0.00705) and high (KW test, p = 0.0763) tagmentation concentrations in comparison to the low one, suggesting that the higher Tn5 concentration is able to recover more genomic fragments when the single cell is insufficiently lysed. The absence of protease had an overall negative effect on the quality. In contrast, libraries built from gDNA or crude cell lysate produced high-quality copy number profiles (data not shown). Based on this observation, we speculated that the lysis step using the new lysis buffer was not sufficient to expose the DNA for efficient tagmentation and we sought to investigate extended lysis times (2 hours and overnight at 4 °C) over a range of protease concentrations. For all experimental conditions the extended lysis improved copy-number quality significantly compared to the shorter lysis (Figure 2c iii-iv; KW tests, p = 5.88753e-30). In addition, we found that increasing the protease concentration beyond the amount used in the microfluidic device had little impact on library quality (Figure S2e; KW test, p = 0.008).

Key to the robust utilization of the approach, we determined that an overnight lysis at 4 °C in combination with the lowest protease concentration provided the best overall performance across all Tn5 concentrations (Figure 2d; mean QS for 2.2nL Tn5 = 0.814 ± 0.367; 3.5nL Tn5 = 0.842 ± 0.318, 6.5nL Tn5 = 0.676 ± 0.451).

For the lysis expansion comparisons (2 hour and 19 hour cold lysis) with the original MF-DLP data (Figure 2d we evaluated how the genome-wide coverage uniformity of our merged DLP+ libraries compared to the MF-DLP dataset. For each condition in the bootstrap analysis, we plotted one Lorenz curve for the merged genome with median coverage breadth and found that merged DLP+ genomes achieved comparable coverage uniformity to the MF-DLP libraries (Figure S2b). It has previously been shown that the MF-DLP single-cell libraries achieved equivalent coverage breadth and uniformity to a standard Nextera bulk genome of equivalent depth (Zahn et al., 2017). It can, therefore, be reasoned that the merged DLP+ genomes also have a comparable quality with that of a bulk library at the same depth. Combined, these results demonstrate that either DLP+ lysis condition sufficiently disrupts cell membranes and proteins and provides adequate access to the genomic DNA during library preparation, generating single-cell libraries with uniformity equivalent to microfluidic DLP.

While cell lysis is critical, we also explored whether adjustments in Tn5 concentration from the MF-DLP range were required. We observed an improvement in library quality with increased Tn5 concentration for insufficiently lysed cells (Figure 2c iii), however, we also detected a significant increase in GC-bias with the increase of Tn5 concentration (Figure 2e). GC-bias is a library characteristic that reflects the correlation between coverage depth of a specific genomic location and its GC-content. Strong GC-bias introduced during library preparation can lead to the under-representation of some genomic regions and over-representation of others. This not only complicates CN inference, it can also result in the dropout of AT- and GC-rich regions due to low coverage, thereby undermining SNV and breakpoint inference in merged clonal or bulk-equivalent genomes (Benjamini and Speed, 2012). The other reaction parameter affecting genome representation was the number of post-tagmentation indexing PCR cycles. We prepared DLP+ libraries using a range of Tn5 concentrations and PCR cycles (Figure 2e, f) and observed an increase of the average GC-content in our single-cell libraries with increasing Tn5 concentration (8 PCR cycles with 2.2 nL Tn5 (2.2nL/8PCR) = 47.1%; 3.5nL Tn5 (3.5nL/8PCR) = 52.7%; 6.5nL Tn5 (6.5nL/8PCR) = 61.5%). In contrast, the increased number of PCR cycles had a very small effect on the GC-content (11 PCR cycles with 2.2 nL Tn5 (2.2nL/11PCR) = 49.4%, (Figure 2e).

To determine if the increased GC bias compromised our ability to generate a high-coverage merged genome, we carried out bootstrap sampling and merging of single-cell libraries from the different experimental conditions and compared the results to our microfluidic GM18507 DLP dataset (Zahn et al., 2017). It should be noted that the MF-DLP dataset was previously shown to produce equivalent breadth and uniformity to that of a bulk genome at the same coverage depth (Zahn et al., 2017). Before the bootstrap analysis, all single-cell libraries were downsampled to achieve the same mean coverage depth of 0.05X (KW test, p= 0.16). Merging 64 downsampled single-cell libraries resulted in 3.16X median coverage depth and 90% median coverage breadth across all libraries (Figure 2f). However, the 6.5nL/8PCR condition and the 2.2nL/11PCR condition had a significant lower genome coverage breadth (86.2% and 89.9%, respectively) compared to the MF-DLP dataset and both the 2.2nL/8PCRand 3.5nL/8PCR conditions (90.6%, 91.3%, 90.7%, respectively; KW test, p = 1.542e-13; a post-hoc Dunn’s test with Benjamini-Hochberg correction showed all comparisons between MF-DLP, 2.2nL/8PCR, nL/8PCR and the 6.5nL/8PCR, 2.2nL/11PCR condition were significant, but there was no significant difference between MF-DLP, 2.23.5 nL/8PCR, 3.5 nL/8PCR conditions) at the same coverage depth (KW test, p = 0.6974). Finally, pooling 128 cells at a mean coverage depth of 0.05X per cell resulted in 96.9% coverage breadth at an aggregated depth of 6.35X for the 2.2nL/8PCR protocol. In comparison, the 6.5nL/8PCR condition achieved significantly less genome coverage (93.2%) at the same depth (KW test, p = 0.3302).

This suggests that the higher GC-bias associated with the increased Tn5 concentration indeed reduced genomic breadth. To evaluate genome-wide uniformity, we again plotted Lorenz curves for each condition in the bootstrap analysis (Figure 2d, Figure S2c). We found that the 6.5 nL Tn5 condition is considerably biased, while there was no major difference between the MF-DLP dataset and the 2.2nL and 3.5 nL Tn5 conditions. For the 2.2nL and 3.5 Nl Tn5 conditions, 64 merged single cells achieve a comparable coverage breadth and uniformity as 64 merged single-cell genomes from the microfluidic dataset (Figure 2f)).

In order to investigate possible breakpoints in the protocol, we next investigated the effects of storing isolated single cells and nuclei on the open-well device. After dispensing and imaging, chips were sealed and stored at −20 °C for 2 months or overnight (”fresh” controls) (Figure S2e). The median library quality was similar for nuclei regardless of storage time, but cells stored for longer had a poorer median QS (Figure S2e. 55% (n = 15/27) of cells stored for 1 day had QS <0.75%, 48% (n = 14/29) of cells stored for 63 days had QS <0.75%, 59% (n = 27/46) of nuclei stored for 1 day had QS <0.75%, 65%(n = 69/106) nuclei stored for 63 days had QS <0.75%, which indicates a similar rate of successful cells/nuclei regardless of storage time.

Finally, we examined the impact of dead cells on library quality. We used imaging information to split the lysis time optimized DLP+ libraries (libraries with overnight lysis, 2.2nL/8PCR and 3.5 nL/8PCR conditions) by cell state (live vs. dead; Figure 2b) and found a significant improvement in the quality of live cells over dead cells (KW test, p = 1.043E-12). Dead cells had a high dropout rate (n= 69% of dead wells failed to produce a library, i.e. less that 250,000 total mapped reads), whereas live cells had a low dropout rate (n= 6% of live wells failed to produced libraries). It is, however, noteworthy that we are able to detect some high-quality CN-profiles in the dead population. This may be related to our inability to distinguish between early and late apoptotic cells with the vital dye staining approache.

## Supplementary Figures

**Figure S 1.**
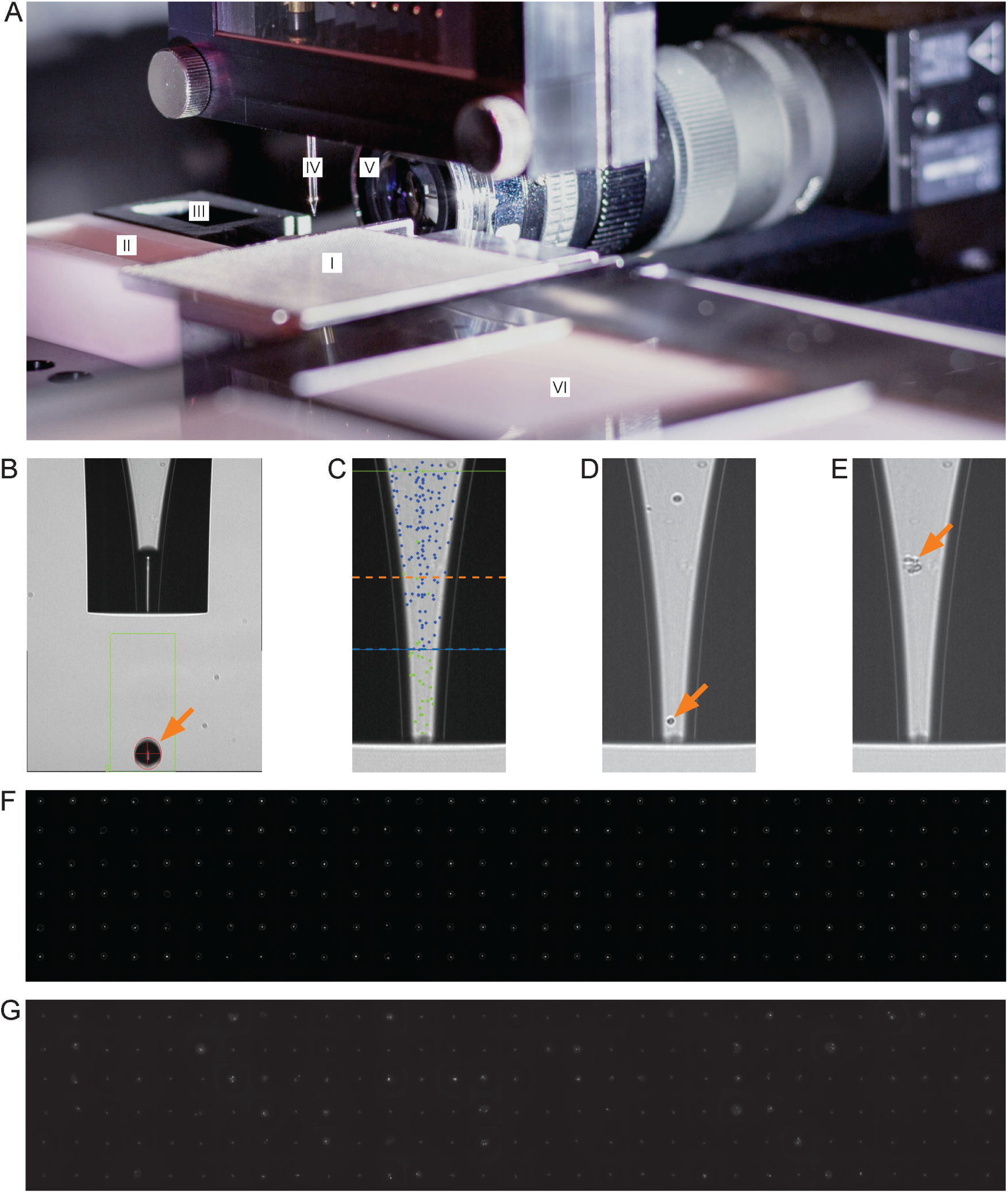
Spotter setup and single-cell isolation relating to method details. a Spotting robot setup featuring: (I) nanowell open-array chip located on customized chip-holder, (II) wash-solution reservoir, (III) active fresh-water wash station, (IV) dispensing nozzle, (V) droplet camera, (VI) chilled target holder. b Brightfield image of the dispensing nozzle. Orange arrow highlights ejected droplet which can range from 300-550 pL in size depending on instrument settings. c Overlay of a brightfield image showing the dispensing nozzle and the mapping density of detected cells. Green dots indicate ejected cells; blue dots indicate cells that were again detected after ejecting a single droplet; dotted blue line shows boundary of cell ejection area/volume; dotted orange line indicates sedimentation boundary. d Automated imaging permits the identification of single cells and target deposition into a nanowell. Cells were deposited if a single cell was detected in the ejection area and no particle was present in the sedimentation area. Orange arrow highlights selected single cell for deposition. e Brightfield image showing contaminating debris (orange arrow). f Montage of 186 fluorescent images of isolated single cells in the bottom of a nanowell using the cellenONE software. Images are aligned according to the array layout. g Montage of 186 fluorescent images from cells dispensed in a limiting dilution. Images are aligned according to the array layout.

**Figure S 2.**
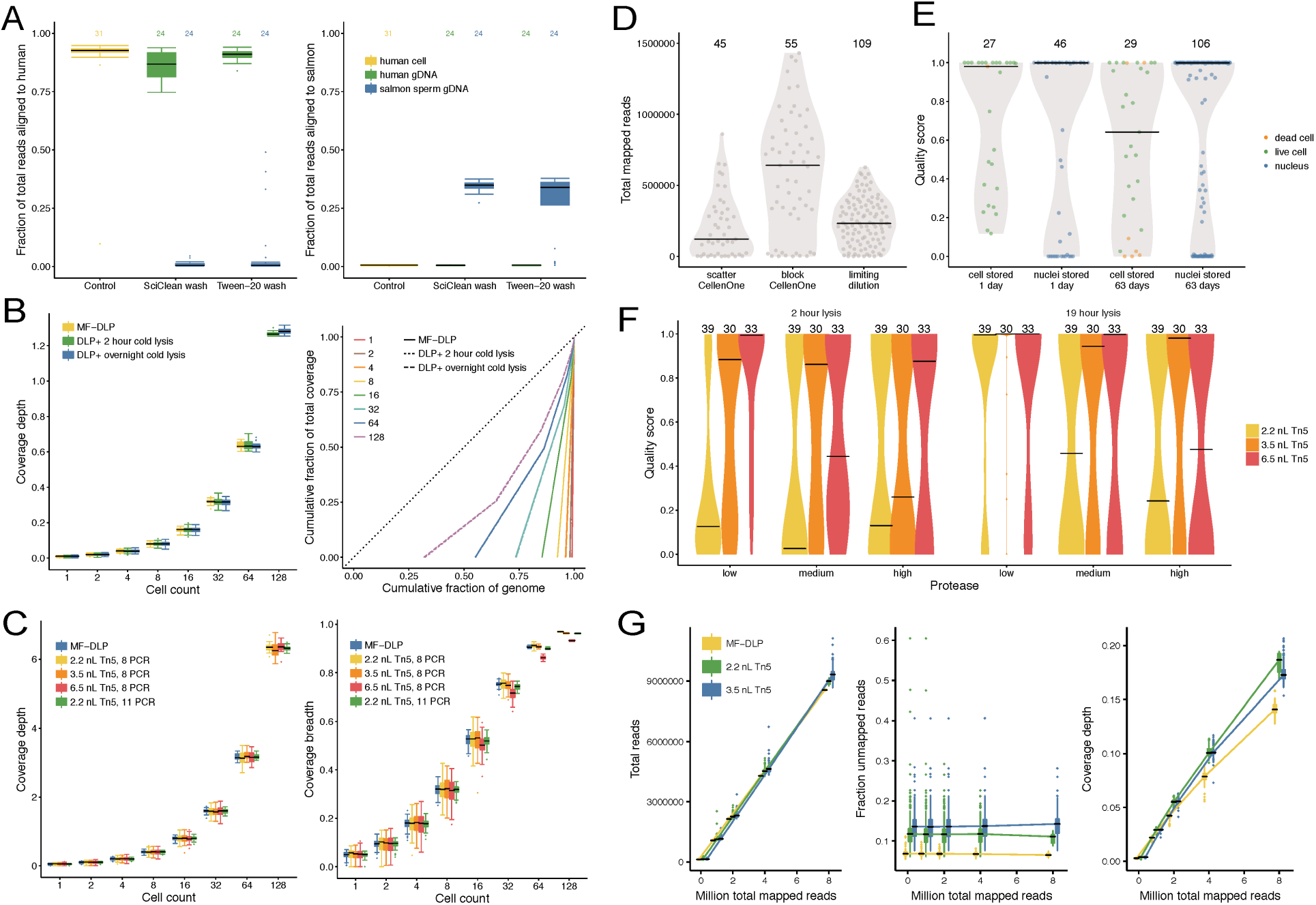
Additional optimization data relating to Figure 2. **a** Cross contamination test for new primer wash routine. A less aggressive wash routine using Tween-20 was tested to replace the primer spotting wash routine using Sciclean as aggressive repetitive Sciclean washes were stripping the nozzle coating and causing issues with drop stability. The new Tween-20 routine was compared to the Sciclean washes by alternating spotting of human and salmon sperm gDNA with a wash step in between and then checking for cross contaminating reads to demonstrate the new wash routine acceptably prevented carryover between wells. The control is human cells dispensed into wells to show typical levels of expected human and salmon reads when primers were dispensed using the Sciclean wash routine. Box plots of fraction of total reads aligned to human and salmon show median and quartiles, the whiskers show the remaining distribution, and dots indicate outliers. b Relating to Figure 2d, the effect of lysis time on coverage breadth of merged single-cell genomes. Bootstrap sampling of single-cell GM18507 libraries prepared using a 2 hour and overnight cold lysis conditions; DLP+ 2 hours (n = 148), DLP+ overnight (n = 133), MF-DLP Zahn et al. (2017) (n = 122). Single-cell libraries were downsampled to a similar median coverage depth. Box plots show median and quartiles, the whiskers show the remaining distribution, and dots indicate outliers. Lorenz curves shows coverage uniformity for merged single-cell genomes. Curves are median merged genomes. Experimental condition and number of merged cells are indicated in the plot. Dotted black line indicates perfectly uniform genome. c Effect of Tn5 concentration and PCR cycles time on coverage breadth of merged single-cell genomes. Bootstrap sampling of single-cell GM18507 libraries prepared using a range of Tn5 concentrations and PCR indexing cycles on the open-array and compared to the MF-DLP dataset (7); DLP+ 2.2 nl Tn5, 8 PCR (n =188), 3.5 nl Tn5, 8 PCR (n = 190), 6.5 nl Tn5, 8 PCR (n = 197), 2.2 nl Tn5, 11 PCR (n = 198), and MF-DLP (7) (n = 122). Single-cell libraries were downsampled to a similar median mean coverage depth. Coverage depth and coverage breadth are shown in boxplots. d Effect of cell dispensing method on total mapped reads, with active selection (cellenONE, spotted in a block of wells or a scatter pattern) or passive limiting dilution dispensing. Black lines show median. e Effect of on-chip storage of isolated cells and nuclei. Quality score of cells and nuclei that were dispensed into nanowells and stored either overnight or for 63 days prior to lysis and library construction. Dots represent individual cells/nuclei. Black lines show median. f Effect of protease concentration on cells. Quality scores of single-cell libraries built with a low, medium, or high concentration of protease in the lysis buffer and lysed for either 2 or 19 hours, followed by library construction with a range of protease concentrations. g Relating to Figure 2g, h comparisons between MF-DLP and DLP+ with two Tn5 concentrations, boxplots show additional sequencing metrics (total reads, fraction unmapped reads, coverage depth). Box plots show median and quartiles, the whiskers show the remaining distribution, and dots indicate outliers. The top column labels state the numbers of cells per condition. Colors represent experimental condition.

**Figure S 3.**
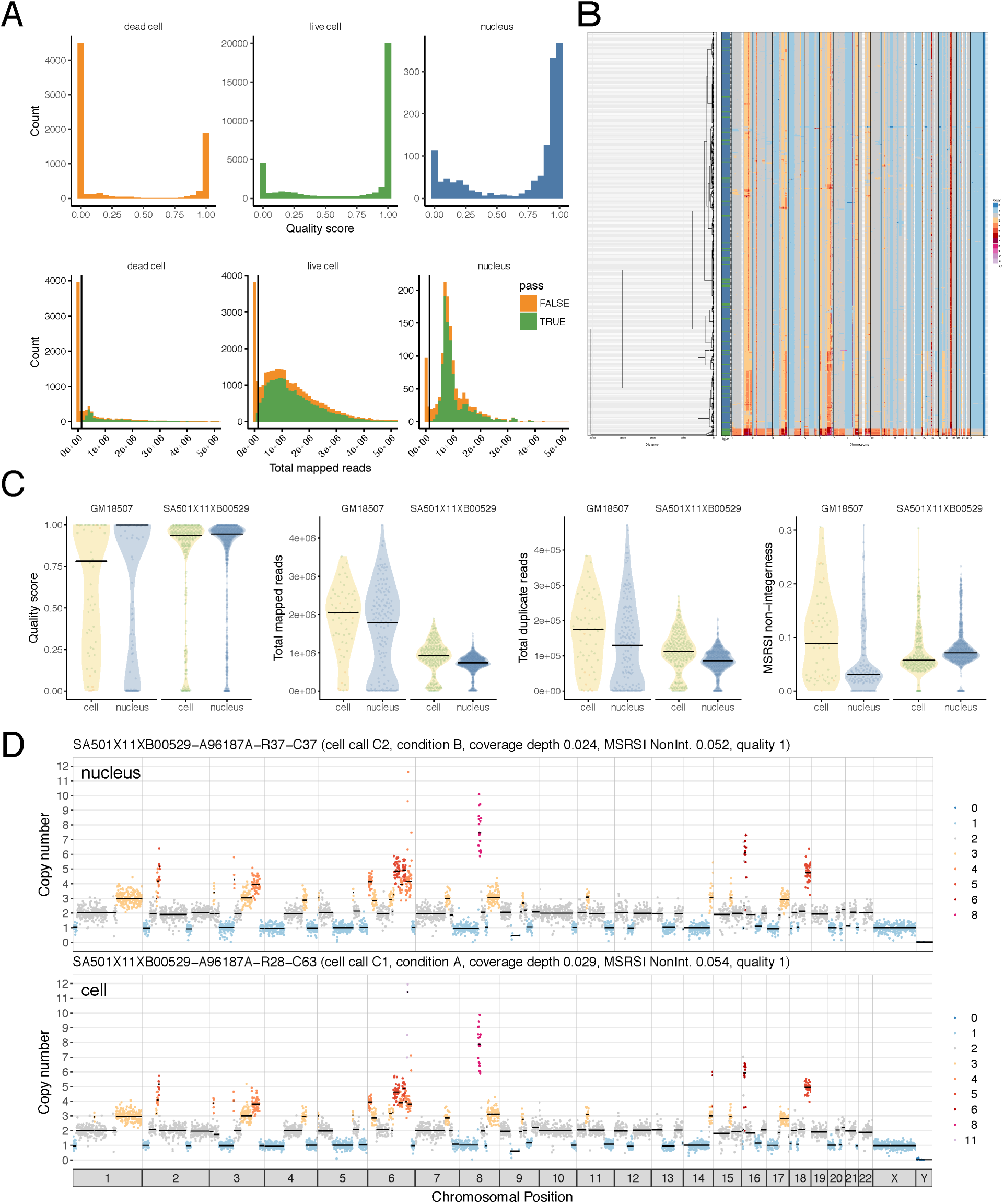
DLP+ produces high quality libraries from cells and nuclei, while dead cells drop out with low read count, relating to Figure 3. **a** Quality score distribution of optimized single-cell libraries, split by dead cells, live cells, and nuclei shows live cells and nuclei have a similar distribution, while dead cells have lower quality. Total mapped reads distribution (orange is cells with QS less than 0.75, and green is cells with QS higher than 0.75), cells with low read counts have low quality score, vertical line represents 125000 reads. b Heatmap of copy number profiles from cells and nuclei shows that cells (green in side bar) and nuclei (blue) cluster together using hierarchical clustering. c Sequencing metrics of single-cell and single-nucleus libraries produced from the same samples. d Example copy number profile from a nucleus and a cell derived from the same sample showing the same copy number clone type.0

**Figure S 4.**
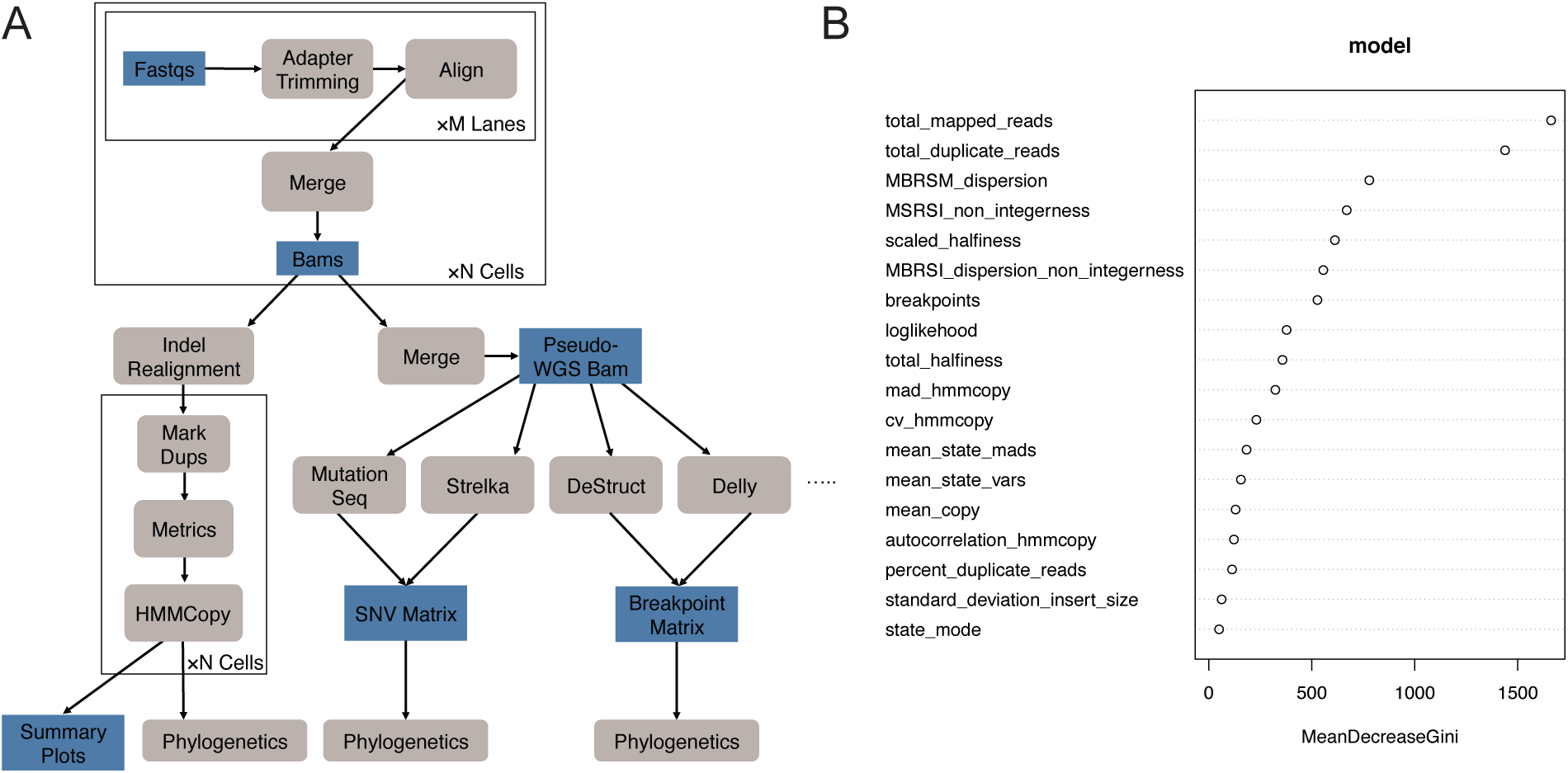
Analysis work flow and classifier feature ranking relating to methods for quantification and statistical analysis. **a** Analysis workflow for alignment and pseudobulk analysis. **b** Random forest classifier feature importance, total mapped reads is of highest importance. Definitions of the features are in methods.

**Figure S 5.**
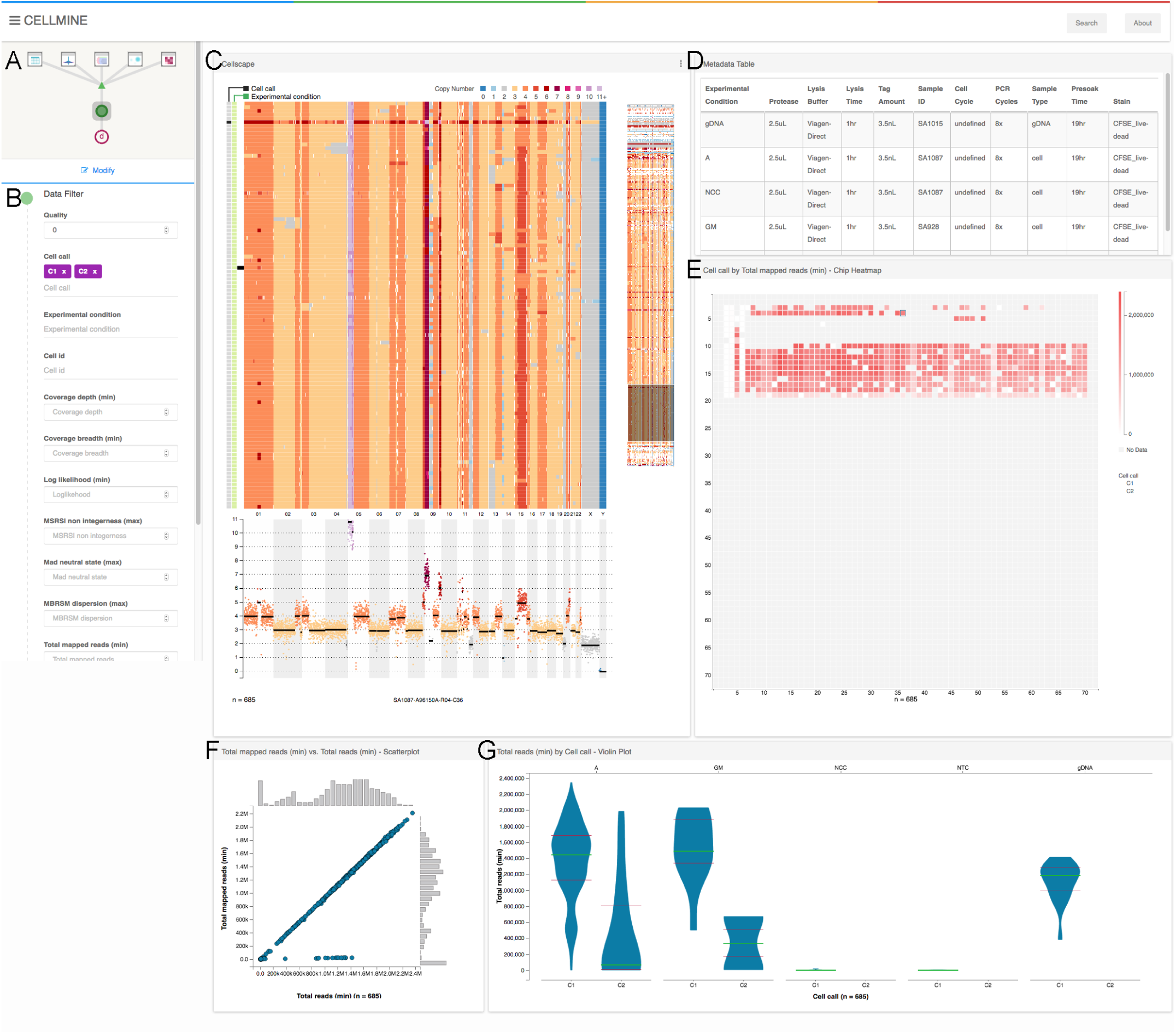
Montage single-cell data visualization relating to methods for Montage for single cell visualization. **a** Interactive parameter selection for the plots and access to data filters. **b** Data filters which can be modified by the user. **c** Cellscape (Smith et al., 2017), a custom heatmap visualization of single-cell copy number profiles. Subpanels include a heatmap detail view, a full heatmap with an interactive slider to select the region of interest in the detail view, and a single-cell copy number profile shown when the user hovers over or selects a cell in the heatmap detail. Side bar shows cell call (C1 indicates live, C2 indicates dead) and experimental condition as colours. **d** Metadata table for interpretation of experimental condition in other plots. **e** Chip heatmap of the open array chip shows a user-selected metric as a colour intensity across wells of the nanowell chip. Each cell of the matrix corresponds to a single well on the chip. **f** Scatter plot shows user selected metrics where a dot is a single-cell genome. Histograms indicate the frequency of values along each axis. **g** Violin plots shows user-selected sequencing metrics and conditions. Lines indicate the median (green) and first and third quantiles (red).

**Figure S 6.**
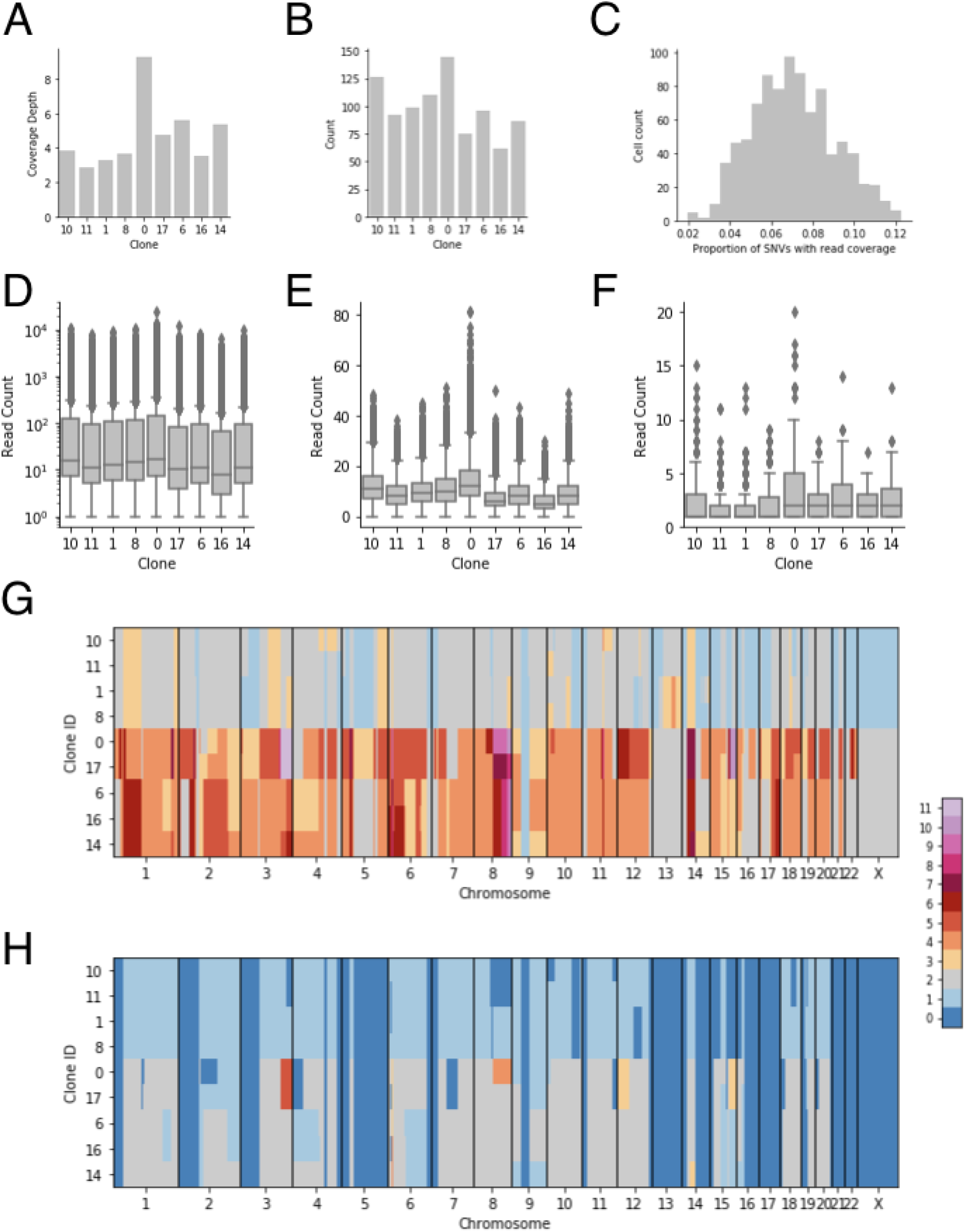
Pseudo-bulk supplementary analysis relating to Figure 4. **a** Coverage in reads reference nucleotide for OV2295 clones. **b** Cell count for OV2295 clones. **c** Histogram of the proportion of SNVs with 1 or more covering reads across cells. **d** Distribution of log read counts per haplotype block as box plots for OV2295 clones. **e** Distribution of read counts per SNV as box plots for OV2295 clones. **f** Distribution of unique read counts per detected breakpoint for OV2295 clones. **g** Total copy number heatmap for each clone (y-axis) across the genome (x-axis). **h** Minor copy number heatmap for each clone (y-axis) across the genome (x-axis).

**Figure S 7.**
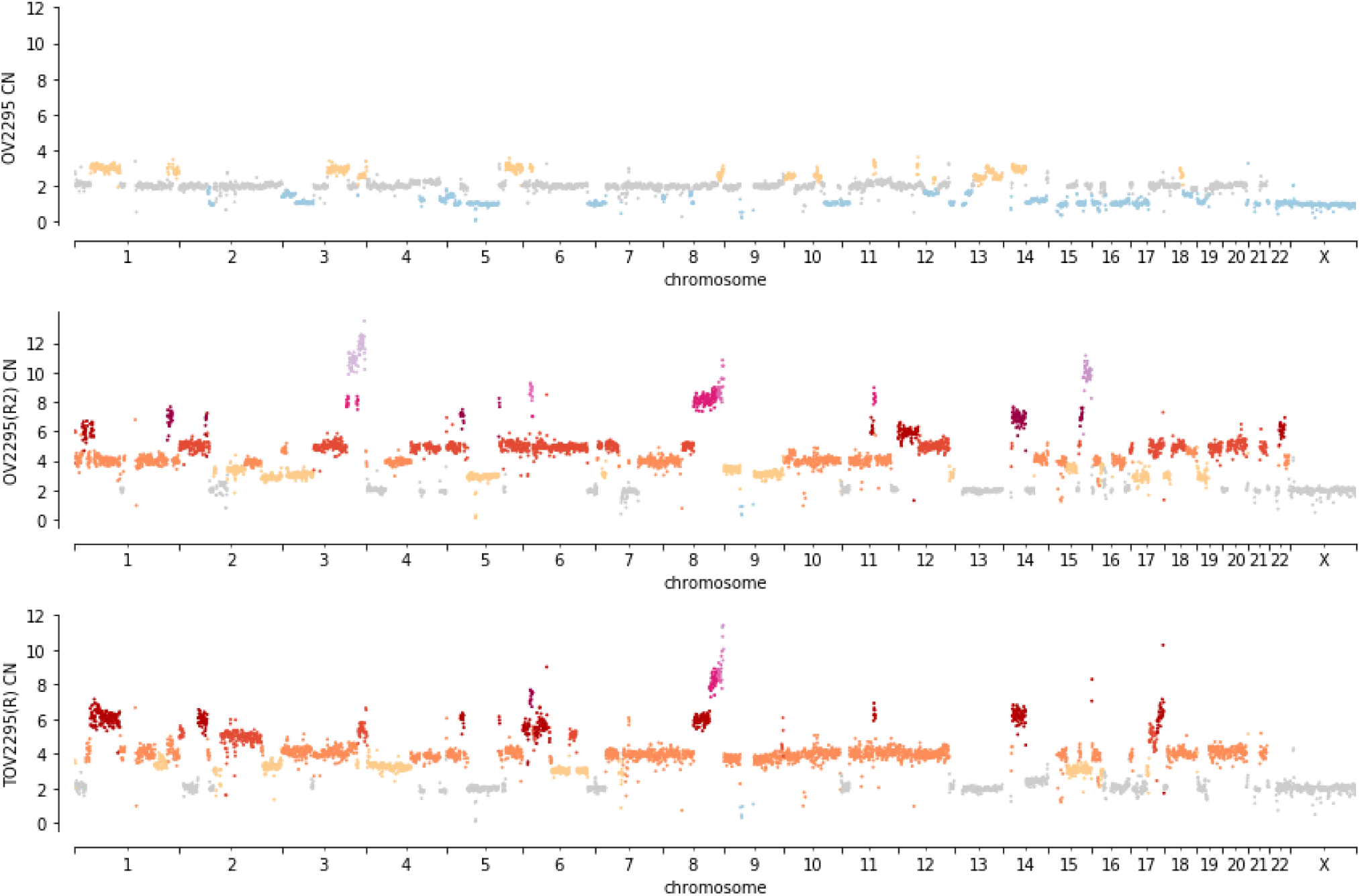
Bulk copy number for each OV2295 sample calculated from DLP+ data.

**Figure S 8.**
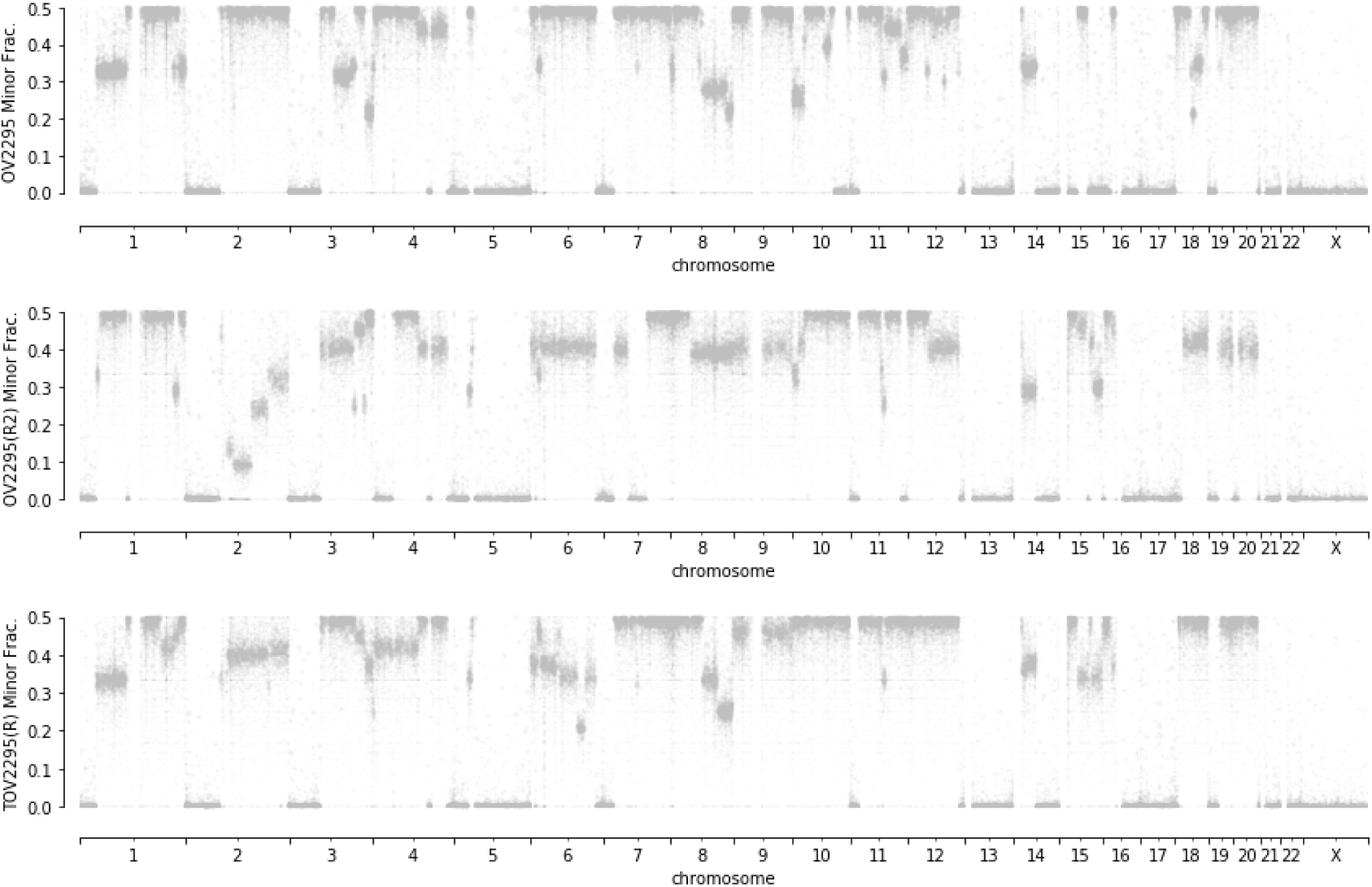
Bulk allele ratios for each OV2295 sample calculated from DLP+ data.

**Figure S 9.**
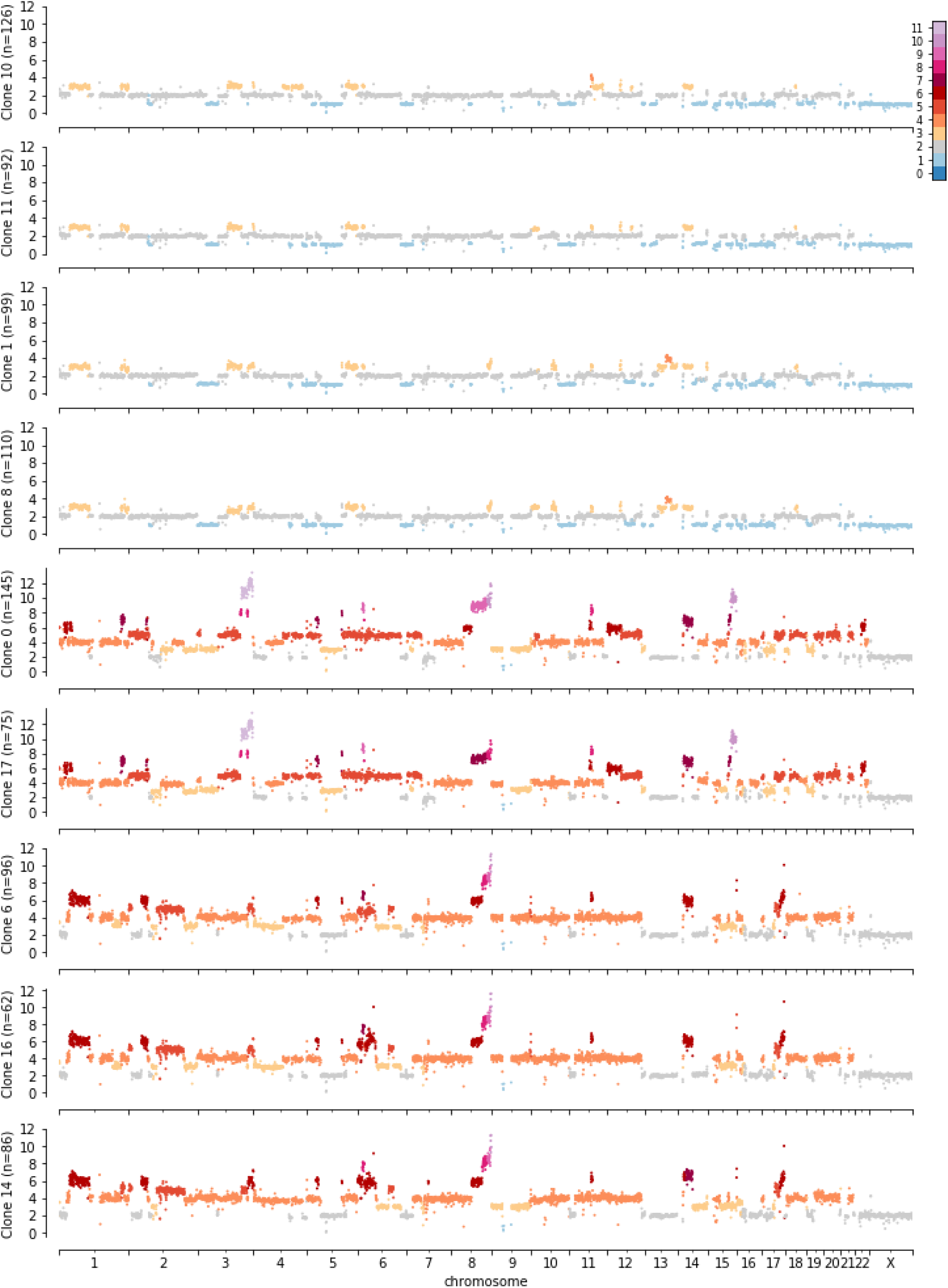
Clone copy number for each OV2295 sample calculated from DLP+ data.

**Figure S 10.**
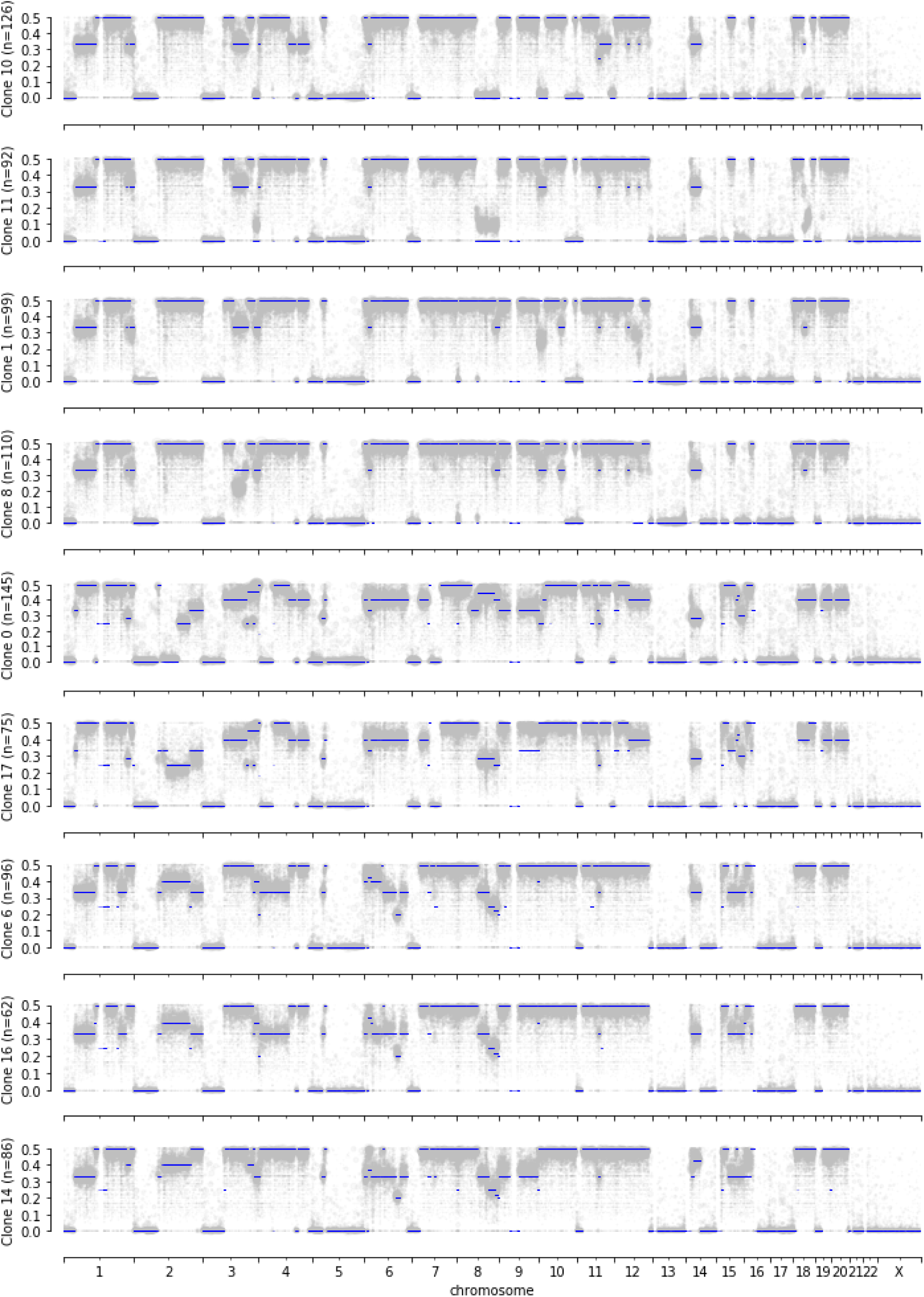
Clone allele ratios for each OV2295 sample calculated from DLP+ data.

**Figure S 11.**
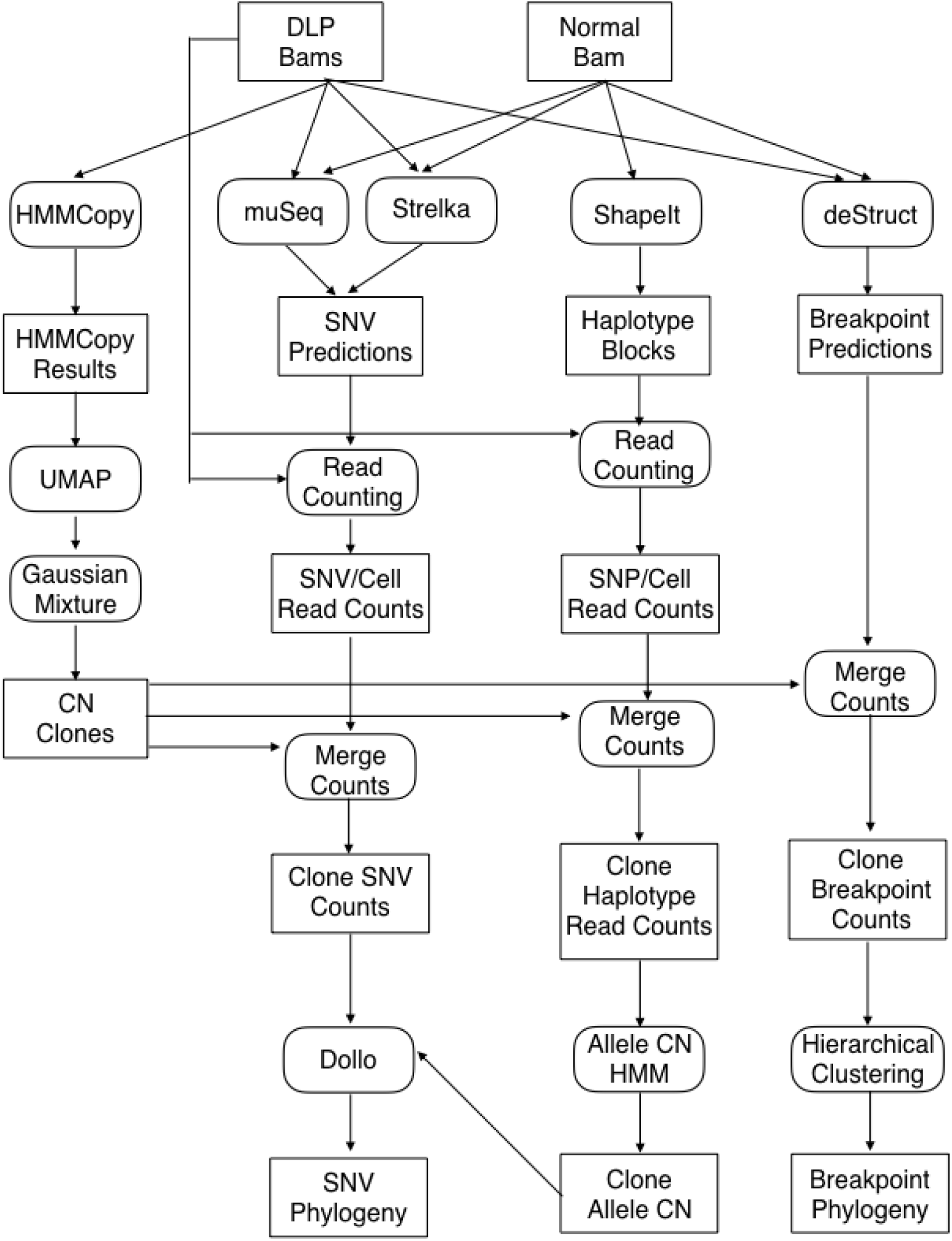
Pseudo-bulk analytical workflow relating to Figure 4. Data (rectangles) and analyses (rounded) for the Pseudo-bulk analytical workflow

**Figure S 12.**
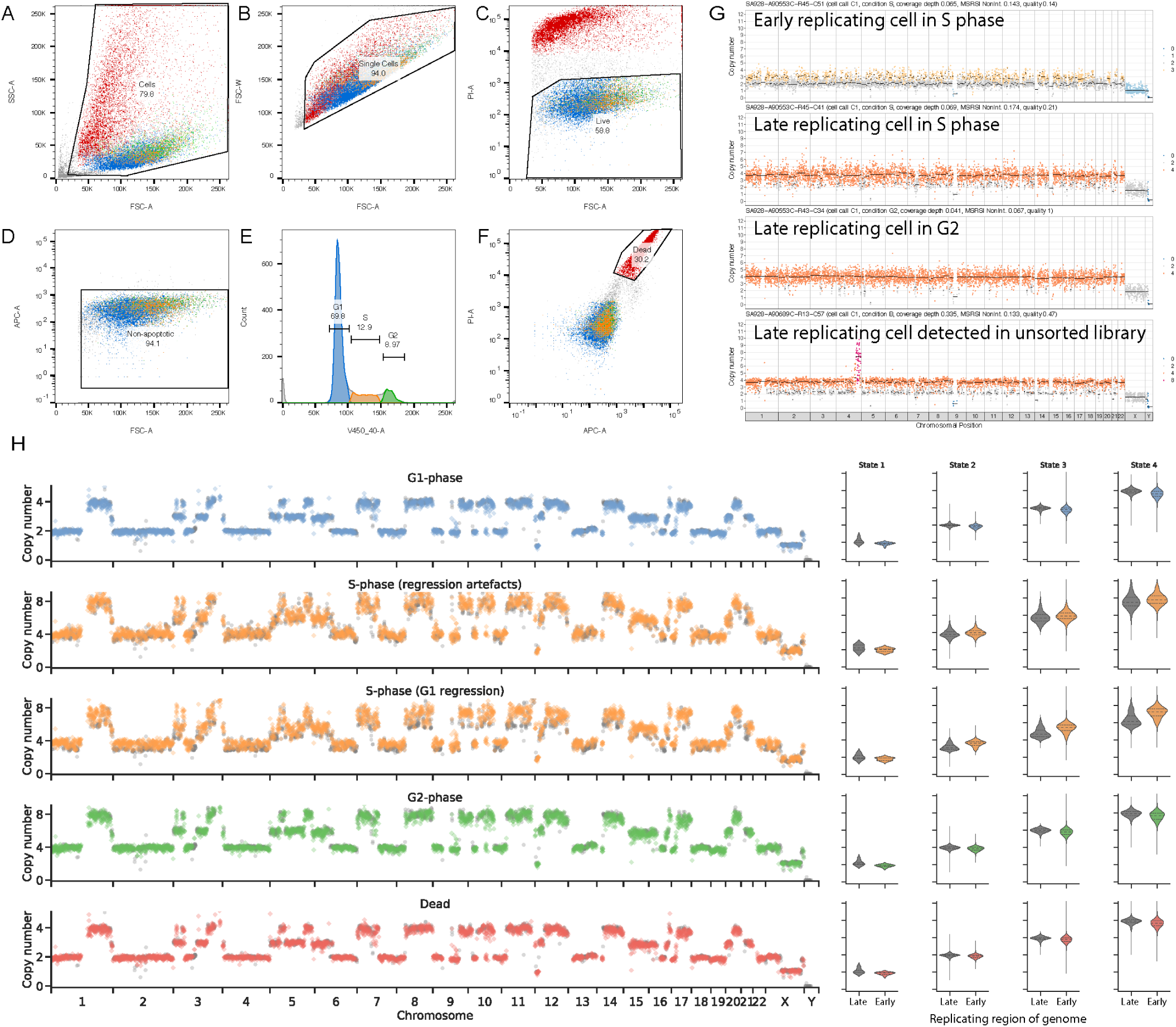
Flow sort gating for cell cycle analysis of G1, S, G2 phase and dead cells by DLP+. **a** Gate for cells. Side scatter area (SSC) vs forward scatter area (FSC) is used to gate out debris (grey) but not dead cells (red) because we will sort them. **b** Gate for single cells. On the cell gate in a, we can use FSC width vs FSC area to gate out doublets if needed for single-cell sorting in a plate. c Gate for live cells. On the gate in **b**, we use PI vs FSC to capture the live cells which are PI low. **d** Gate for non-apoptotic cells. On the live cell gate in **c**, we use Caspase 3/7 (APC-A vs FSC) to exclude apoptotic cells which are Caspase 3/7 high from our live cell population. **e** Gate for cell cycle phases in live cells. On the live cell gate established in **a-d**, we use the DNA content of the cells measured by Hoechst 33342 staining (V459/40-A)to gate the G1 (blue), S (orange), and G2 (green) phases of the cell cycle. **f** Gate for dead cells. On the gate for single cells established in b, we gate on the PI high, Caspase 3/7 high dead cells (red). **g** Example GM18507 cells in S phase and G2 with early replicating regions leading and late replicating regions lagging, including a cell from an unsorted experiment, showing we can detect these cells without preselecting the population. Colours correspond to integer HMM copy-number states (Ha et al., 2012); black lines indicate segment medians. **h** Read distributions of early and late replicating regions in an aneuploid cell line. Violin plots of read distributions of early and late replicating regions of the genome of the flow sorted T-47D cell line split by cell state.

## Supplementary Tables

**Table S1:** Summarization of the library construction conditions for the cells in this resource.

**Table S2:** DLP+ single cell sequencing metrics per single cell library. Sequencing, alignment and quality metrics for each single cell reported.

**Table S3:** Omnibus table of statistical comparisons of the DLP+. Table sections refer to contrasts of data subgroups, the statistical method and results.

**Table S4:** Summarization of high quality cell sequencing metrics.

**Table S5:** SNV state per clone in OV2295 samples. Columns are ‘chrom’: chromosome, ‘coord’: coordinate of SNV, ‘clone_id’: clone identifier, ‘state’: state of SNV. The SNV state is translated as follows, 0: no reads, 1: absent, 2: heterozygous, 3: homozygous.

**Table S6:** Breakpoint state per clone in OV2295 samples. Columns are ‘chromosome_n’: chromosome of break end n, ‘strand_n’: orientation of break end n, ‘position_n’: position of break end n, ‘clone_id’: clone identifier, ‘state’: state of breakpoint. The breakpoint state is translated as follows, 0: absent, 1: present.

**Table S7:** Allele specific copy number state per clone in OV2295 samples. Columns are ‘chr’: chromosome of bin, ‘start’: start of bin, ‘end’: end of bin, ‘total_cn’, ‘minor_cn’, ‘major_cn’: total, minor and major copy number of bin in the given clone, ‘clone_id’: clone identifier.

**Table S8:** Results of UMAP and Gaussian Mixture clustering by copy number of OV2295 samples. Columns are ‘cell_id’: cell identifier, ‘gm_umap_cluster’: cluster index as clone identifier, ‘umap1’, ‘umap2’: first, second component of UMAP dimensionality reduction.

## References

Baslan, T., Kendall, J., Rodgers, L., Cox, H., Riggs, M., Stepansky, A., Troge, J., Ravi, K., Esposito, D., Lakshmi, B. et al. (2012). Genome-wide copy number analysis of single cells. Nature Protocols 7, 1024–1041.

Benjamini, Y. and Speed, T. P. (2012). Summarizing and correcting the GC content bias in high-throughput sequencing. Nucleic Acids Research 40, e72.

Burleigh, A., McKinney, S., Brimhall, J., Yap, D., Eirew, P., Poon, S., Ng, V., Wan, A., Prentice, L., Annab, L. et al. (2015). A co-culture genome-wide RNAi screen with mammary epithelial cells reveals transmembrane signals required for growth and differentiation. Breast Cancer Research 17, 4.

Delaneau, O., Marchini, J. and Zagury, J.-F. (2011). A linear complexity phasing method for thousands of genomes. Nat Methods 9, 179–81.

Ding, J., Bashashati, A., Roth, A., Oloumi, A., Tse, K., Zeng, T., Haffari, G., Hirst, M., Marra, M. A., Condon, A. et al. (2012). Feature-based classifiers for somatic mutation detection in tumour-normal paired sequencing data. Bioinformatics 28, 167–75.

Eirew, P., Steif, A., Khattra, J., Ha, G., Yap, D., Farahani, H., Gelmon, K., Chia, S., Mar, C., Wan, A. et al. (2015). Dynamics of genomic clones in breast cancer patient xenografts at single-cell resolution. Nature 518, 422–426.

Gao, R., Kim, C., Sei, E., Foukakis, T., Crosetto, N., Chan, L.-K., Srinivasan, M., Zhang, H., Meric-Bernstam, F. and Navin, N. (2017). Nanogrid single-nucleus RNA sequencing reveals phenotypic diversity in breast cancer. Nature Communications 8, 228.

Gawad, C., Koh, W. and Quake, S. R. (2014). Dissecting the clonal origins of childhood acute lymphoblastic leukemia by single-cell genomics. Proceedings of the National Academy of Sciences 111, 17947–17952.

Gawad, C., Koh, W. and Quake, S. R. (2016). Single-cell genome sequencing: current state of the science. Nature Reviews Genetics 17, 175 EP-.

Goldstein, L. D., Chen, Y.-J. J., Dunne, J., Mir, A., Hubschle, H., Guillory, J., Yuan, W., Zhang, J., Stinson, J., Jaiswal, B. et al. (2017). Massively parallel nanowell-based single-cell gene expression profiling. BMC genomics 18, 519.

Ha, G., Roth, A., Khattra, J., Ho, J., Yap, D., Prentice, L. M., Melnyk, N., McPherson, A., Bashashati, A., Laks, E. et al. (2014). TITAN: inference of copy number architectures in clonal cell populations from tumor whole-genome sequence data. Genome Res. 24, 1881–93.

Ha, G., Roth, A., Lai, D., Bashashati, A., Ding, J., Goya, R., Giuliany, R., Rosner, J., Oloumi, A., Shumansky, K. et al. (2012). Integrative analysis of genome-wide loss of heterozygosity and monoallelic expression at nucleotide resolution reveals disrupted pathways in triple-negative breast cancer. Genome research 22, 1995–2007.

Haldar, M., Hancock, J. D., Coffin, C. M., Lessnick, S. L. and Capecchi, M. R. (2007). A conditional mouse model of synovial sarcoma: insights into a myogenic origin. Cancer Cell 11, 375–388.

Hansen, R. S., Thomas, S., Sandstrom, R., Canfield, T. K., Thurman, R. E., Weaver, M., Dorschner, M. O., Gartler, S. M. and Stamatoyannopoulos, J. A. (2010). Sequencing newly replicated DNA reveals widespread plasticity in human replication timing. Proceedings of the National Academy of Sciences 107, 139–144.

Hou, Y., Song, L., Zhu, P., Zhang, B., Tao, Y., Xu, X., Li, F., Wu, K., Liang, J., Shao, D. et al. (2012). Single-Cell Exome Sequencing and Monoclonal Evolution of a JAK2-Negative Myeloproliferative Neoplasm. Cell 148, 873–885.

Knouse, K. A., Lopez, K. E., Bachofner, M. and Amon, A. (2018). Chromosome Segregation Fidelity in Epithelia Requires Tissue Architecture. Cell 0.

Knouse, K. A., Wu, J., Whittaker, C. A. and Amon, A. (2014). Single cell sequencing reveals low levels of aneuploidy across mammalian tissues. Proc Natl Acad Sci U S A 111, 13409–14.

L6tourneau, I. J., Quinn, M. C. J., Wang, L.-L., Portelance, L., Caceres, K. Y., Cyr, L., Delvoye, N., Meunier, L., de Ladurantaye, M., Shen, Z. et al. (2012). Derivation and characterization of matched cell lines from primary and recurrent serous ovarian cancer. BMC cancer 12, 379.

Leung, K., Klaus, A., Lin, B. K., Laks, E., Biele, J., Lai, D., Bashashati, A., Huang, Y.-F., Aniba, R., Moksa, M. et al. (2016). Robust high-performance nanoliter-volume single-cell multiple displacement amplification on planar substrates. Proceedings of the National Academy of Sciences 113, 8484–8489.

Lohr, J. G., Adalsteinsson, V. A., Cibulskis, K., Choudhury, A. D., Rosenberg, M., Cruz-Gordillo, P., Francis, J. M., Zhang, C.-Z., Shalek, A. K., Satija, R. et al. (2014). Whole-exome sequencing of circulating tumor cells provides a window into metastatic prostate cancer. Nature Biotechnology 32,479.

Macosko, E. Z., Basu, A., Satija, R., Nemesh, J., Shekhar, K., Goldman, M., Tirosh, I., Bialas, A. R., Kamitaki, N., Martersteck, E. M. et al. (2015). Highly Parallel Genome-wide Expression Profiling of Individual Cells Using Nanoliter Droplets. Cell 161, 1202–1214.

McInnes, L. and Healy, J. (2018). UMAP: Uniform Manifold Approximation and Projection for Dimension Reduction.

McPherson, A., Roth, A., Laks, E., Masud, T., Bashashati, A., Zhang, A. W., Ha, G., Biele, J., Yap, D., Wan, A. et al. (2016). Divergent modes of clonal spread and intraperitoneal mixing in high-grade serous ovarian cancer. Nat Genet 48, 758–67.

McPherson, A., Shah, S. P. and Sahinalp, S. C. (2017a). deStruct: Accurate Rearrangement Detection using Breakpoint Specific Realignment.

McPherson, A. W., Roth, A., Ha, G., Chauve, C., Steif, A., de Souza, C. P. E., Eirew, P., Bouchard-Côté, A., Aparicio, S., Sahinalp, S. C. et al. (2017b). ReMixT: clone-specific genomic structure estimation in cancer. Genome Biol 18, 140.

Navin, N., Kendall, J., Troge, J., Andrews, P., Rodgers, L., McIndoo, J., Cook, K., Stepansky, A., Levy, D., Esposito, D. et al. (2011). Tumour evolution inferred by single-cell sequencing. Nature 472, 90.

Ni, X., Zhuo, M., Su, Z., Duan, J., Gao, Y., Wang, Z., Zong, C., Bai, H., Chapman, A. R., Zhao, J. et al. (2013). Reproducible copy number variation patterns among single circulating tumor cells of lung cancer patients. Proceedings of the National Academy of Sciences 110, 21083–21088.

Sanders, A. D., Falconer, E., Hills, M., Spierings, D. C. J. and Lansdorp, P. M. (2017). Single-cell template strand sequencing by Strand-seq enables the characterization of individual homologs. Nature Protocols 12, 1151–1176.

Saunders, C. T., Wong, W. S. W., Swamy, S., Becq, J., Murray, L. J. and Cheetham, R. K. (2012). Strelka: accurate somatic small-variant calling from sequenced tumor-normal sample pairs. Bioinformatics 28, 1811–7.

Smith, M. A., Nielsen, C. B., Chan, F. C., McPherson, A., Roth, A., Farahani, H., Machev, D., Steif, A. and Shah, S. P. (2017). E-scape: interactive visualization of single-cell phylogenetics and cancer evolution. Nature Methods 14,549 EP-.

Soto, M., Raaijmakers, J. A., Bakker, B., Spierings, D. C. J., Lansdorp, P. M., Foijer, F. and Medema, R. H. (2017). p53 Prohibits Propagation of Chromosome Segregation Errors that Produce Structural Aneuploidies. Cell Rep 19,2423–2431.

The International HapMap Consortium, T. I. H. (2005). A haplotype map of the human genome. Nature 437, 12991320.

Vitak, S. A., Torkenczy, K. A., Rosenkrantz, J. L., Fields, A. J., Christiansen, L., Wong, M. H., Carbone, L., Steemers, F. J. and Adey, A. (2017). Sequencing thousands of single-cell genomes with combinatorial indexing. Nature Methods 14, 302–308.

Wang, Y., Waters, J., Leung, M. L., Unruh, A., Roh, W., Shi, X., Chen, K., Scheet, P., Vattathil, S., Liang, H. et al. (2014). Clonal evolution in breast cancer revealed by single nucleus genome sequencing. Nature 512, 155.

Woodfine, K., Fiegler, H., Beare, D. M., Collins, J. E., McCann, O. T., Young, B. D., Debernardi, S., Mott, R., Dunham, I. and Carter, N. P. (2004). Replication timing of the human genome. Human Molecular Genetics 13,191–202.

Wu, L., Zhang, X., Zhao, Z., Wang, L., Li, B., Li, G., Dean, M., Yu, Q., Wang, Y., Lin, X. et al. (2015). Full-length single-cell RNA-seq applied to a viral human cancer: applications to HPV expression and splicing analysis in HeLa S3 cells. GigaScience 4, 51.

Zahn, H., Steif, A., Laks, E., Eirew, P., VanInsberghe, M., Shah, S. P., Aparicio, S. and Hansen, C. L. (2017). Scalable whole-genome single-cell library preparation without preamplification. Nature Methods 14, 167–173.

Ziegenhain, C., Vieth, B., Parekh, S., Reinius, B., Guillaumet-Adkins, A., Smets, M., Leonhardt, H., Heyn, H., Hellmann, I. and Enard, W. (2017). Comparative Analysis of Single-Cell RNA Sequencing Methods. Molecular Cell 65, 631-643.e4.

Zong, C., Lu, S., Chapman, A. R. and Xie, X. S. (2012). Genome-Wide Detection of Single-Nucleotide and Copy-Number Variations of a Single Human Cell. Science 338, 1622–1626.

